# Negative density dependence promotes persistence of a globally rare yet locally abundant plant species (*Oenothera coloradensis*)

**DOI:** 10.1101/2023.10.25.563615

**Authors:** Alice E. Stears, Bonnie Heidel, Maria Paniw, Roberto Salguero-Gómez, Daniel C. Laughlin

## Abstract

1. Identifying the mechanisms underlying the persistence of rare species has long been a motivating question for ecologists. Classical theory implies that community dynamics should be driven by common species, and that natural selection should not allow small populations of rare species to persist. Yet, a majority of the species found on Earth are rare. Consequently, several mechanisms have been proposed to explain their persistence, including negative density dependence, demographic compensation, vital rate buffering, asynchronous responses of sub-populations to environmental heterogeneity, and fine-scale source-sink dynamics. Persistence of seeds in a seed bank, which is often ignored in models of population dynamics, can also buffer small populations against collapse.
2. We used integral projection models (IPMs) to examine the population dynamics of *Oenothera coloradensis*, a rare, monocarpic perennial forb, and determine whether any of five proposed demographic mechanisms for rare species persistence contribute to the long-term viability of two populations. We also evaluated how including a discrete seed bank stage changed population models for this species.
3. Including a seed bank stage in population models had a significant positive impact on modeled *O. coloradensis* population growth rate. Using IPMs that included a discrete seedbank state, we found that negative density-dependence was the only supported mechanism for the persistence of this rare species.
4. *Synthesis*: IPMs of two populations of the rare species *O. coloradensis* emphasize the importance of including cryptic life stages such as seed banks in demographic models, but fail to provide strong support for most of the proposed mechanisms of rare species persistence. We propose that high micro-site abundance in a spatially heterogeneous environment enables this species to persist, allowing it to sidestep the demographic and genetic challenges of small population size that rare species typically face. These results emphasize that globally rare species can employ many different strategies for persistence, including the somewhat counter-intuitive phenomenon of local abundance. This reinforces the need for customized management and conservation strategies that mirror the diversity of mechanisms that allow rare species persistence.

## Introduction

Determining how and why populations of rare species persist has been a goal for ecologists since the discipline’s inception (Levins & Culver, 1971; Drury, 1974). Theoretically, low population size is a final step on a trajectory toward extinction (Stanley, 1979) or the first step toward ubiquity (Spear, Walsh, Ricciardi, & Vander Zanden, 2021). Yet, small but stable populations of rare species exist in every ecosystem and taxonomic group (Magurran & Henderson, 2011). In fact, a large proportion of species globally – as many as 35% of plant species, for example— can be considered naturally rare (Enquist et al., 2019). The prevalence of rarity suggests it is an evolutionarily stable strategy rather than a stop along the path toward extinction or invasion, and implies that there must be both fundamental and realized niches that are available for rare species to occupy. A growing body of evidence demonstrates the importance of rare species for biological processes, including their impacts on community stability (Arnoldi, Loreau, & Haegeman, 2019), and functional composition (Burner et al., 2022), which in turn impact ecosystem function (Lyons, Brigham, Traut, & Schwartz, 2005).

Effective conservation and management of rare species require an understanding of both the conditions causing rarity initially, and the mechanisms that allow rare species to persist. Causes of rarity can vary from highly-specific habitat requirements (Sgarbi & Melo, 2018), to adverse impacts of anthropogenic environmental change (Vincent, Bornand, Kempel, & Fischer, 2020). In order to then persist in a state of rarity, a species must overcome any of multiple potential challenges, primarily the negative effects of demographic, environmental, and genetic stochasticity, defined as random variation in vital rates (e.g., survival, reproduction), abiotic conditions, or genetic allele frequencies (May, 1973). Stochastic deleterious events can cause extirpation or even extinction of rare species, since there may not be enough unaffected individuals or subpopulations to “rescue” the affected population (Nei, Maruyama, & Chakraborty, 1975). Rare species that maintain populations over time typically do so by employing demographic strategies that compensate for the adverse effects of small population size. There are five main strategies that allow persistence of rare populations (Fig. 1) (Dibner, Peterson, Louthan, & Doak, 2019): negative density-dependence (Rovere & Fox, 2019), demographic compensation (Villellas, Doak, García, & Morris, 2015), vital rate buffering (Pfister, 1998; Hilde et al., 2020), asynchronous responses between subpopulations (Abbott, Doak, & Peterson, 2017), and fine-scale source-sink dynamics (Kauffman, Pollock, & Walton, 2004; Pulliam, 1988). Negative density-dependence occurs when the growth rate (*λ*) of a population increases at small population size. Demographic compensation occurs when different vital rates are affected in opposing ways by the same perturbation in the environment, which can help maintain a relatively constant population *λ* in response to environmental variation. Vital rate buffering occurs when the variability of vital rates decreases as the vital rate becomes more important(i.e., has a higher elasticity), which prevents negative effects of temporal variation on the *λ* across time (Tuljapurkar, 1989). Spatial asynchrony occurs when subpopulations close to one another have different or even opposing growth rates, resulting in a stable population-wide *λ*. Fine-scale source-sink dynamics occur when there is gene flow between subpopulations that bolsters the size and genetic diversity of very small subpopulations, which again results in a stable population-level *λ*. Each of these mechanisms can act independently, but also can interact or overlap (Dibner et al., 2019).

Here, we identify which of these mechanisms contribute to the persistence or population growth of a rare, endemic plant species, *Oenothera coloradensis* (Rydberg) W.L. Wagner & Hoch (Onagraceae). We use integral projection models (IPMs) (Easterling, Ellner, & Dixon, 2000) that include a discrete seed bank population state. IPMs are flexible models of population dynamics that are constructed using regression models that describe vital rate change across a continuous state variable such as size. IPMs have multiple advantages including better performance with small datasets than traditional matrix models (Ramula, Rees, & Buckley, 2009), and direct incorporation of covariates of interest directly into vital rate models. We built these models with two objectives in mind.

Our first objective was to determine if including information about the seed bank significantly altered population models for *O. coloradensis*. Seed banks can serve as important reservoirs of genetic diversity and buffer populations against collapse (Vitalis, Glémin, & Olivieri, 2004), and can be critical for monocarpic perennials such as *O. coloradensis* that only flower once in their lifetime (Rees et al., 2006). For these reasons, we expected that a soil seed bank is important for *O. coloradensis*, and that its inclusion in IPMs would increase population *λ*. Seed banks are often not included in population models because their parameters can be very difficult to estimate, but previous work shows that including them can significantly alter model outcomes (Paniw, Quintana-Ascencio, Ojeda, & Salguero-Gómez, 2017; Nguyen, Buckley, Salguero-Gómez, & Wardle, 2019).

Our second objective was to identify whether any of the five aforementioned persistence mechanisms was acting to maintain *O. coloradensis* populations. This species occurs in habitats that naturally experience frequent, highly localized disturbance, meaning that some subpopulations might be negatively affected by flood, for example, while other nearby subpopulations are simultaneously thriving due to lack of disturbance. Additionally, previous matrix population models constructed for this species in the 1990s found substantial variation in *λ* across space and time (Floyd & Ranker, 1998). The population-wide pattern of asynchronous habitat disturbance also could make source-sink dynamics important. Finally, we have evidence of large fluctuations in the number of plants within subpopulations (Heidel, Tuthill, & Wallace, 2021), which suggest that *λ* decreases at high population size and increases at low population size. Therefore, we predicted that density dependence, small-scale source-sink dynamics and asynchronous responses between subpopulations would be important mechanisms of persistence for *O. coloradensis*.

**Figure 1.**
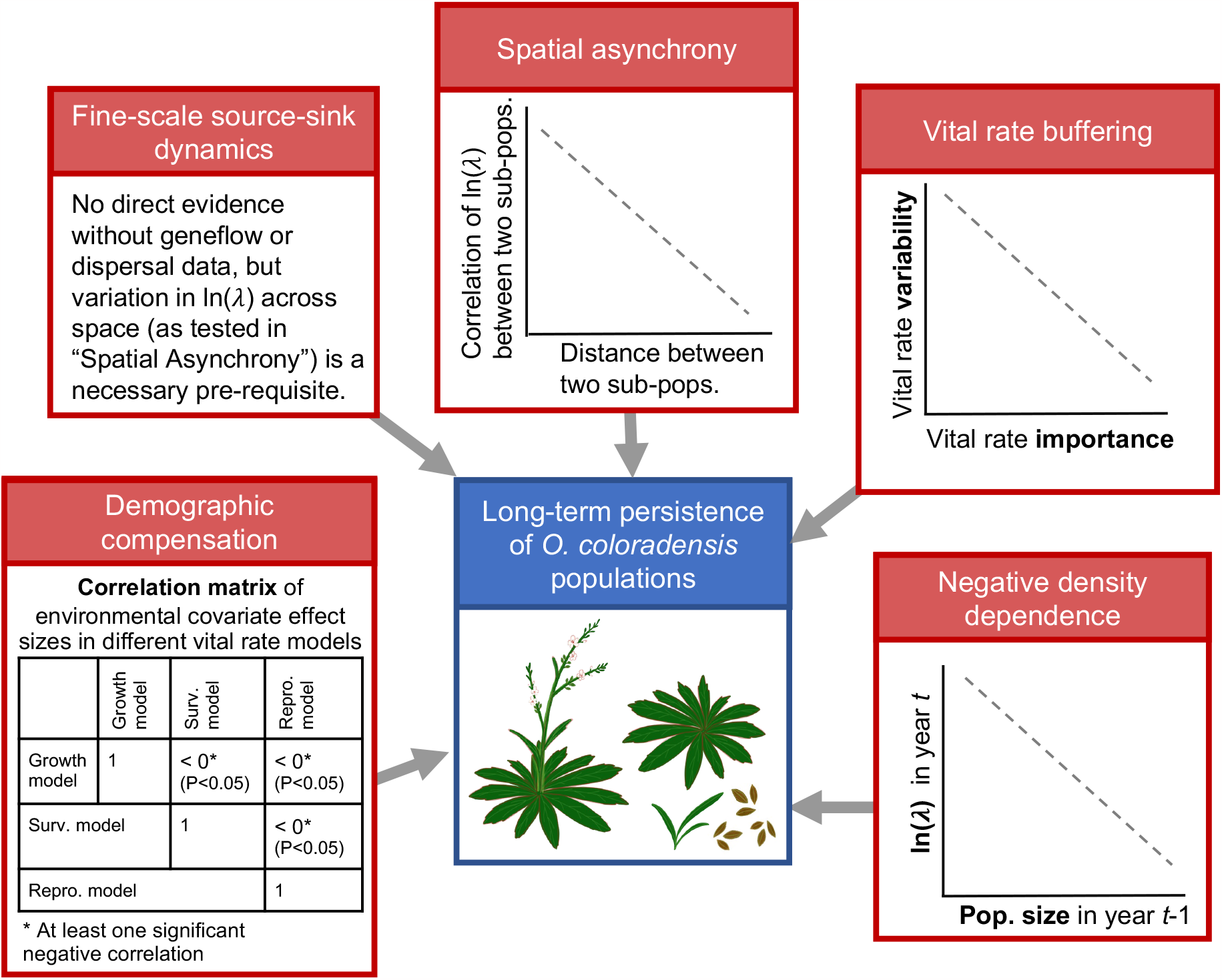
Evidence that would be required to support each of the five mechanisms that can contribute to long-term viability of small populations of rare species.

## Materials and Methods

### Species Description

*Oenothera coloradensis* (Onagraceae) (Wagner, Krakos, & Hoch, 2013) is an herbaceous, monocarpic perennial plant that occurs in frequently disturbed, mesic or wet meadows, and riparian floodplains (Fertig, 2000). Non-reproductive individuals consist of a rosette of leaves with a fleshy taproot. Flowering typically occurs after several years, when individuals bolt and produce a 10-30 cm long floral stalk. Individuals typically die after reproducing–93% of the time in populations we observed. Frequent disturbance such as flooding that reduces competing species and removes litter is important, especially for successful seedling recruitment (Fertig, 2000). All known *O. coloradensis* populations lie within a 7,000-hectare area that includes southeast Wyoming, northern Colorado, and a small part of southwest Nebraska (Fig. S1). The only known population on Federal land occurs on the F. E. Warren Air Force Base near Cheyenne, WY (FEWAFB). The Soapstone Prairie Natural Area (Soapstone), owned by the city of Fort Collins, CO, has the largest known population of *O. coloradensis* individuals (Heidel et al., 2021). The U.S. Fish and Wildlife Service (USFWS) designated *O. coloradensis* as a “threatened” species under the Endangered Species Act in 2000 (Endangered and Threatened Wildlife and Plants, 2000). Managers were concerned that habitat loss due would lead to extinction of this naturally rare species. However, based on subsequent monitoring, the USFWS determined that the species does not appear to be on a trajectory toward extinction, but fluctuates naturally. As a result, O. coloradensis was de-listed in 2019. As a result, *O. coloradensis* was de-listed in 2019 (Endangered and Threatened Wildlife and Plants, 2019).

Previous work established that *O. coloradensis λ* is particularly impacted by recruitment of seedlings (Floyd & Ranker, 1998). Seed banks are also likely important, since years of high seedling density are not necessarily preceded by years of high rates of flowering and seed production (Heidel et al., 2021). The *O. coloradensis* seed bank has not been studied directly, but a greenhouse seed study showed that an average of 58% of seeds produced by a parent plant are viable, and that a viable seed has a 20% probability of germinating after two months of cold stratification (Burgess, Hild, & Shaw, 2005). These rates did not change meaningfully over five years (see “Supplementary Material: Species Information” for more detail).

### Demographic Data Collection

We conducted a three-year demographic study of *O. coloradensis* across six spatially distinct subpopulations, three in the FEWAFB population (”Unnamed creek”, “Crow creek”, and “Diamond creek”) and three at the Soapstone population (”Meadow”, “HQ3” and “HQ5”)(Table S1; Fig. S1). In early summer 2018, we established three 2x2 m^2^ quadrats in each of these subpopulations, resulting in 18 plots (Table S1). Plants larger than 3 cm are typically “non-seedling” plants at least one year in age. In each study plot, we tagged and mapped each unique “non-seedling” individual and recorded longest leaf length, reproductive status, and seed production for each (see “Supplementary Information: Seed Production Estimation” for more detail). Individuals smaller than 3 cm in leaf length are typically seedlings that germinated that year, occur at high density, and are less likely to survive than non-seedling plants. Due to these factors, we tallied seedlings in each plot, but did not map or tag them. In subsequent 2019 and 2020 censuses, we mapped and tagged new “non-seedling” individuals, and re-measured all surviving individuals from previous years. Sample size in a given year at a subpopulation ranged from 48 to 1527 individuals (Table S1). All mapping, tagging, and leaf measurements took place between late May and early July, during the peak of vegetative growth for this species.

### Environmental Measurements

To determine the effect of temporal variation in climate on *O. coloradensis* populations, we used modeled, population-level temperature and precipitation data from PRISM (PRISM Climate Group; Oregon State University, 2021), which we refer to as “environmental covariates”. We calculated the mean temperature of both the growing season (April in *year*_*t*_ – August in *year*_*t*_) and preceding winter season (September in *year*_*t*−1_ – March in *year*_*t*_) for each year of vital rate data collection at FEWAFB and Soapstone. We also calculated total precipitation for each “water year” (October in *year*_*t*−1_ to September in *year*_*t*_), which we used in place of growing season precipitation because the shortgrass steppe ecosystem in which *O. coloradensis* occurs receives a majority of its annual precipitation in the form of snow, and melting snow from the previous winter likely drives springtime seedling recruitment. Average temperature of the previous winter is also likely important for seedling recruitment, because seed germination is triggered by cold stratification (Burgess et al., 2005). Growing season temperature and precipitation are likely important for growth, survival, and reproductive output of non-seedling plants.

### Vital Rate Models

We used data from the three-year demographic monitoring study detailed above to parameterize models of *O. coloradensis* vital rates (shown in Fig. 2; parameters of fitted vital rate functions are shown in Table S2). We first estimated continuous vital rate functions describing how survival probability, growth, flowering probability, and seed production in *year*_*t*+1_ each vary as a function of longest leaf size in *year*_*t*_. We also estimated a continuous vital rate function describing the distribution of new recruit size in *year*_*t*+1_. Finally, we estimated discrete vital rate parameters describing the probability of seeds produced in *year*_*t*_ either entering the seed bank or germinating in *year*_*t*+1_, as well as the probability of seeds in the seed bank in *year*_*t*_ either staying in the seed bank or germinating in *year*_*t*+1_.

We first created “global” models for each continuous vital rate (Table. 1). We modeled survival probability (*s*(*z*)) as a function of log-transformed leaf size (ln(*size*_*t*_)) using generalized linear models with binomial error distributions. Flowering individuals were excluded from the data used to fit survival models, since *O. coloradensis* is a monocarpic perennial that nearly always dies after flowering. We modeled probability of flowering (*Pb*(*z*)) as a function of using generalized linear models with binomial error distributions, and was predicted by ln(*size*_*t*_) plus ln(*size*_*t*_)-squared. We modeled seed production (*b*(*z*)) as a function of ln(*size*_*t*_) using negative binomial models because the count data was over-dispersed. We only used data from flowering plants to fit these models, using the “glm.nb” function from the “MASS” R package (Venables & Ripley, 2002). We described growth, or the distribution of plant size in *year*_*t*+1_ (*G*(*z*^*′*^, *z*)) as a series of Normal distributions with mean = *μ*_*s*_ and standard deviation = *σ*_*s*_. *μ*_*s*_ was modeled as a function of ln(*size*_*t*_) using a linear model with Gaussian error. *σ*_*s*_ was the residual standard error of this linear model. We described the distribution of recruit size in *year*_*t*+1_ (*c*_*o*_(*z*′)) as a Normal distribution with the mean *μ*_*r*_, and the standard deviation *σ*_*r*_. *μ*_*r*_ and *σ*_*r*_ were the mean and standard deviation of observed longest leaf size in *year*_*t*+1_.

We then used these “global” model structures to fit different versions of continuous vital rate functions, each of which described vital rate processes at different temporal and spatial scales. Models were fit using data from the first transition (2018-2019), the second transition (2019-2020), or pooled across both transitions. We also used data from a single subpopulation, a single population, or pooled across both populations. We additionally fit models that expanded on the “global” model structures by including density dependence terms and/or environmental covariates (water year precipitation, mean annual growing season temperature, or mean annual winter temperature). When density dependence or environmental covariates were included, we used AIC model selection to confirm that including these covariates improved model fit. All continuous vital rate models, regardless of scale, were parameterized using data from “non-seedling plants” as well as seedlings. We incorporated seedlings into the continuous, non-seedling dataset using a process detailed in “Supplementary Information: Seedling Data.”

We estimated discrete vital rates for seeds uniformly across both populations and years, using data from both greenhouse and field-based germination and seed viability studies. (Table 1). We did not have the data required to determine how these rates changed across subpopulations or in response to abiotic variation, due to the difficulties of estimating *in situ* seed germination and death. We used the following parameters to estimate discrete seed vital rate parameters: viable seed germination rate (germ. rate) = 0.16, viability rate of seeds produced by a parent plant (viab. rate) = 0.58, rate of natural seed death in the seed bank (death rate) = 0.10 (see “Supplementary Information: Discrete Vital Rate Parameters” for more detail).

### Population Models

We used estimates of discrete and continuous *O. coloradensis* vital rates to parameterize a suite of integral projection models (IPMs) for *O. coloradensis*. We then used these models to address the objectives outlined in the Introduction.

**Figure 2.**
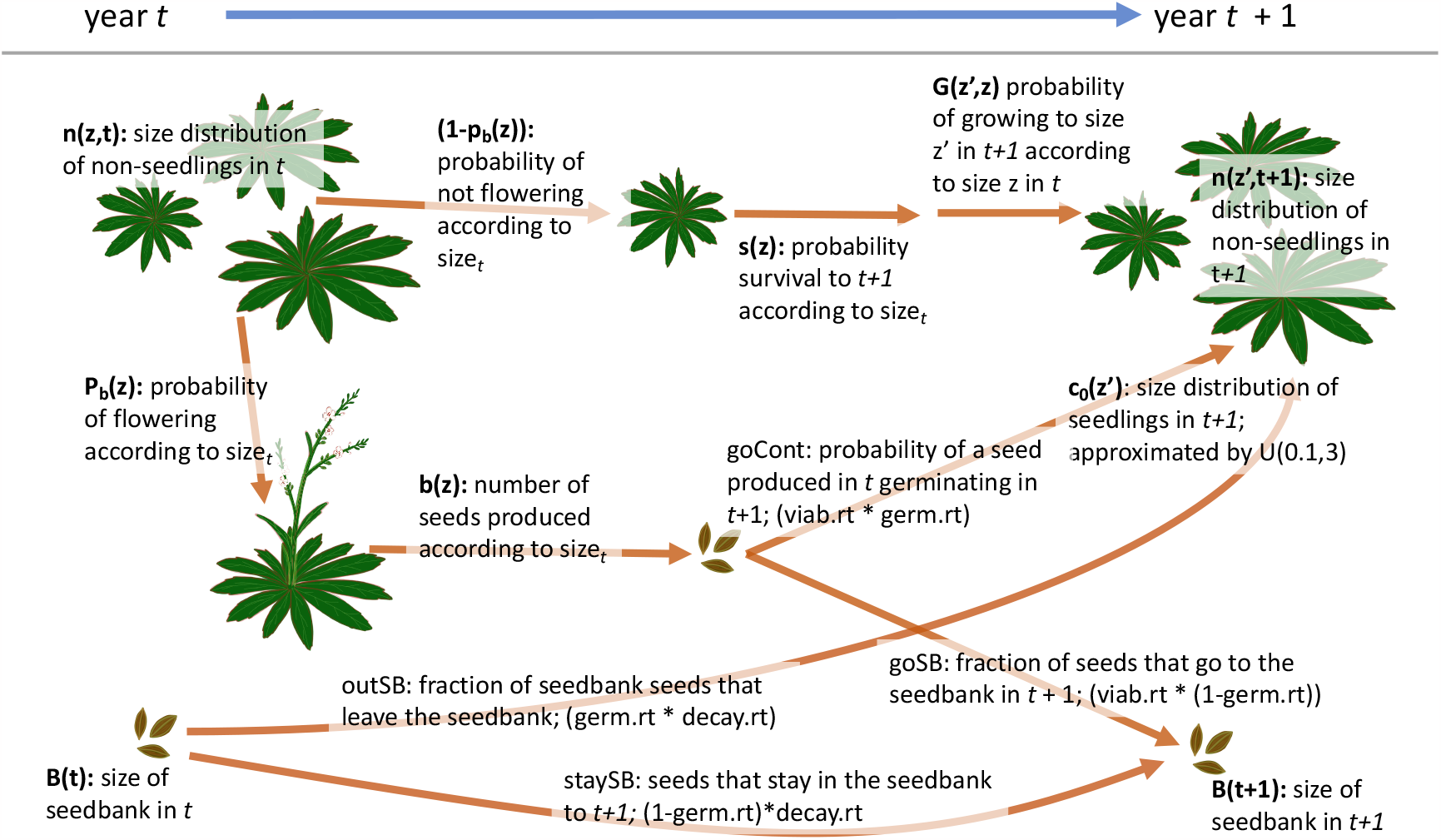
Diagram of the *O. coloradensis* life-cycle, labeled with notation used in vital rate equations shown in Table 1. Based on model structures and notation from: (Paniw et al., 2017; Merow et al., 2014; Ellner et al., 2016). “germ.rt” = germination rate, “viab.rt” = viability rate, “decay.rt” = decay rate.

### Objective 1: Importance of the Seed bank Stage

We used two different IPMs to determine whether explicitly including a discrete seed bank stage in a population model leads to significantly different outcomes relative to a model without a seed bank stage. We first created a density-independent IPM using continuous vital rate functions parameterized with data from both Soapstone and FEWAFB. This model had a single continuous, size-based population state, and did not include a seed bank state (Table 2: IPM “A”; Eqn. 1). This IPM used a kernel structure where the continuous, above-ground population state (*n*(*z*′, *t* + 1)) at time *t* + 1 was described by the following equation:

**Table 1:** Description of vital rates used in *O. coloradensis* IPMs.

| Vital Rate | Description | Model |
| --- | --- | --- |
| $pEstab$ | $P(\text{seed produced in } year_t \text{ establishes as a seedling in } year_{t+1})$ | $pEstab = \frac{\text{Num. new recruits in } year_{t+1}}{\text{Num. seeds produced in } year_t}$ |
| $goCont$ | $P(\text{seed produced in } year_t \text{ germinates in } year_{t+1})$ | $goCont = \text{viab. rate} \times \text{germ. rate}$ |
| $outSB$ | $P(\text{seed bank seed in } year_t \text{ germinates in } year_{t+1})$ | $outSB = \text{germ. rate} \times (1 - \text{death rate})$ |
| $goSB$ | $P(\text{seed produced in } year_t \text{ goes into the seed bank in } year_{t+1})$ | $goSB = \text{viab. rate} \times (1 - \text{germ. rate})$ |
| $staySB$ | $P(\text{seed bank seed in } year_t \text{ stays in the seed bank in } year_{t+1})$ | $staySB = (1 - \text{germ. rate}) \times (1 - \text{death rate})$ |
| $Survival (s(z))$ | $P(\text{survival from } year_t \text{ to } year_{t+1})$ | $\text{logit}(\text{survival}) \sim \beta_0 + \beta_1(\ln(\text{size}_t)) + \epsilon$ |
| $Flowering (Pb(z))$ | $P(\text{flowering in } year_t)$ | $\text{logit}(\text{flowering}) \sim \beta_0 + \beta_1(\ln(\text{size}_t)) + \beta_2(\ln(\text{size}_t)^2) + \epsilon$ |
| $Seed prod. (b(z))$ | Seed production in $year_t$ | $\exp(\text{seed number}) \sim \beta_0 + \beta_1(\ln(\text{size}_t)) + \epsilon$ |
| $Growth (G(z', z))$ | Distribution of $\ln(\text{longest leaf size})$ in $year_t$ | $G(z', z) = N(\mu_s, \sigma_s);$<br>$\mu_s \sim \beta_0 + \beta_1(\ln(\text{size}_t)) + \epsilon;$<br>$\sigma_s \sim RSE(\beta_0 + \beta_1(\ln(\text{size}_t)) + \epsilon)$ |
| $Recruit size (c_o(z'))$ | Distribution of new recruit size in $year_t$ | $c_o(z') = N(\mu_r, \sigma_r);$<br>$\mu_r = \text{mean}(\text{size of recruits in } t);$<br>$\sigma_r = \text{stnd. dev. (size of recruits in } t)$ |

\* RSE = residual standard error

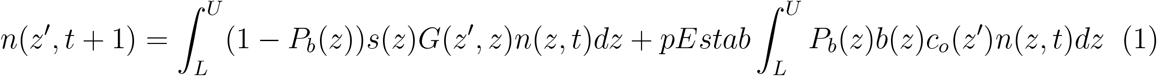

We then created an IPM that included both a continuous, size-based stage for above-ground individuals, and a discrete seed bank state (Table 2: IPM “B”; Eqns. 2 & 3) (Ellner & Rees, 2006; Rees et al., 2006; Paniw et al., 2017). This model used the same continuous vital rate functions as IPM A, but also included discrete values describing the probabilities of seeds produced in *year*_*t*_ germinating or going into the seeds bank in *year*_*t*+1_, and the probabilities of seeds in the seed bank in *year*_*t*_ germinating or persisting in the seed bank in *year*_*t*+1_. This IPM with two population states used a kernel structure where the continuous, above-ground population state (*n*(*z*′, *t* + 1)) and the seed bank state (*B*(*t* + 1)) at time *t* + 1 were described by the following equations:

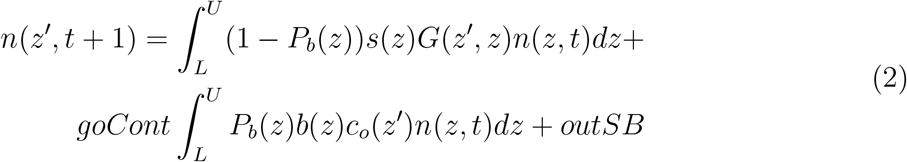

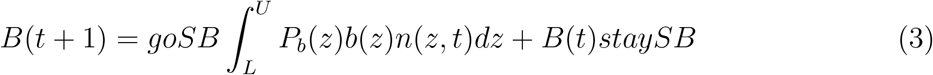

**Table 2:**
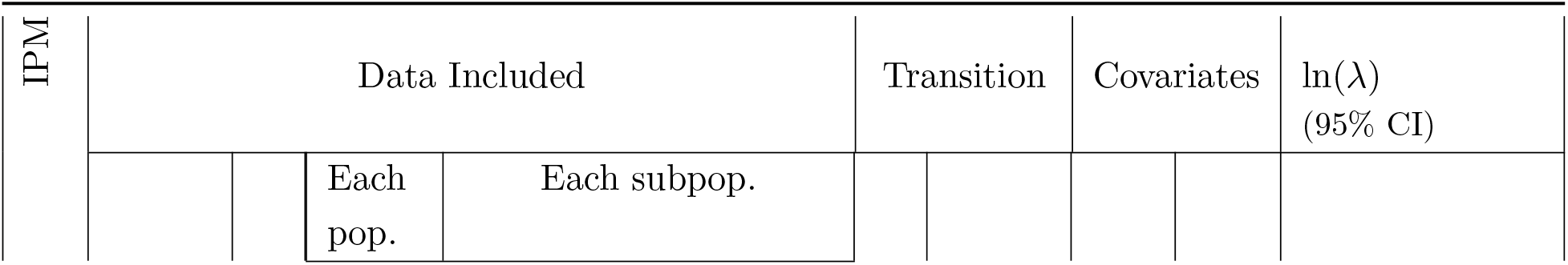
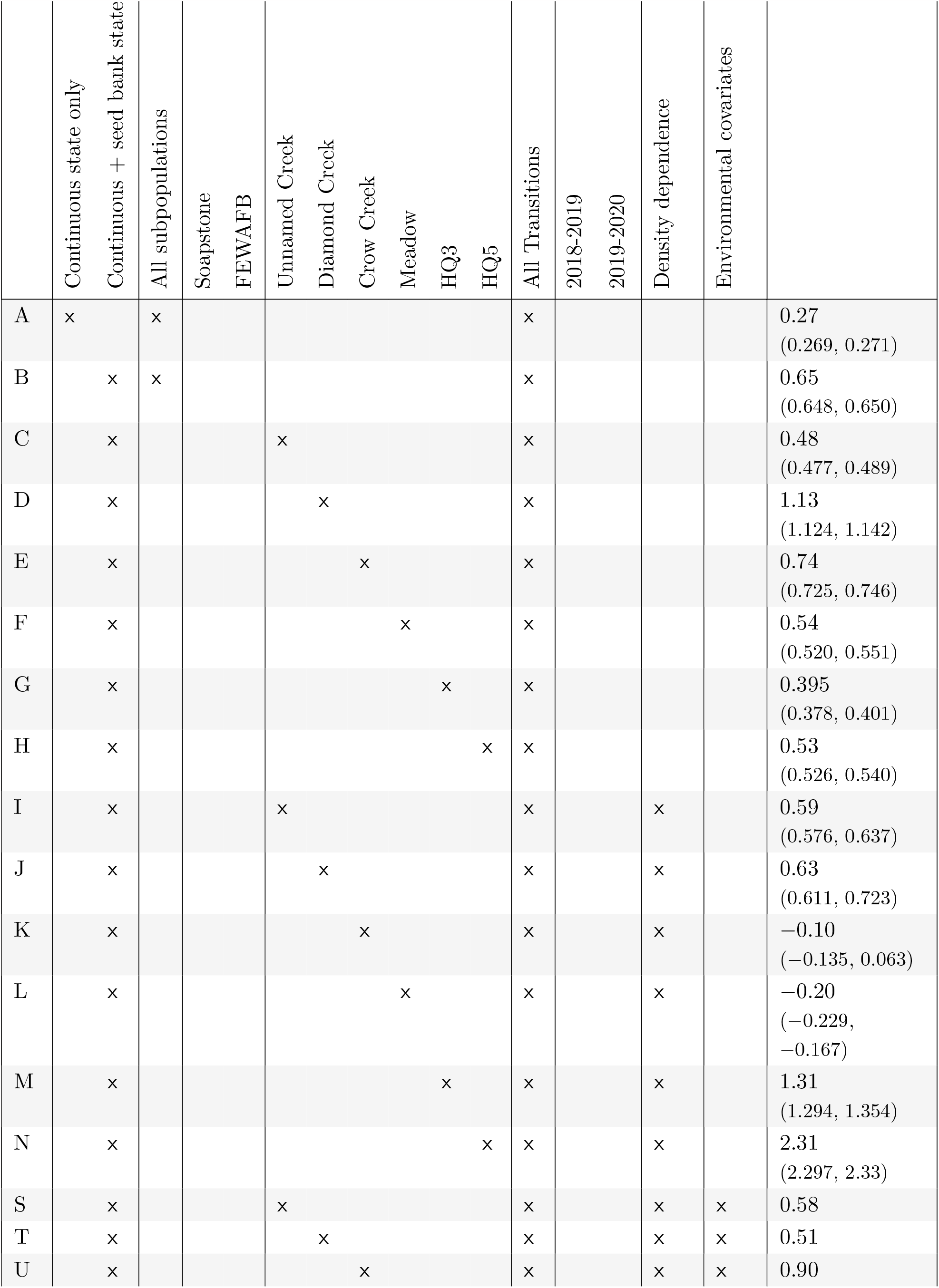
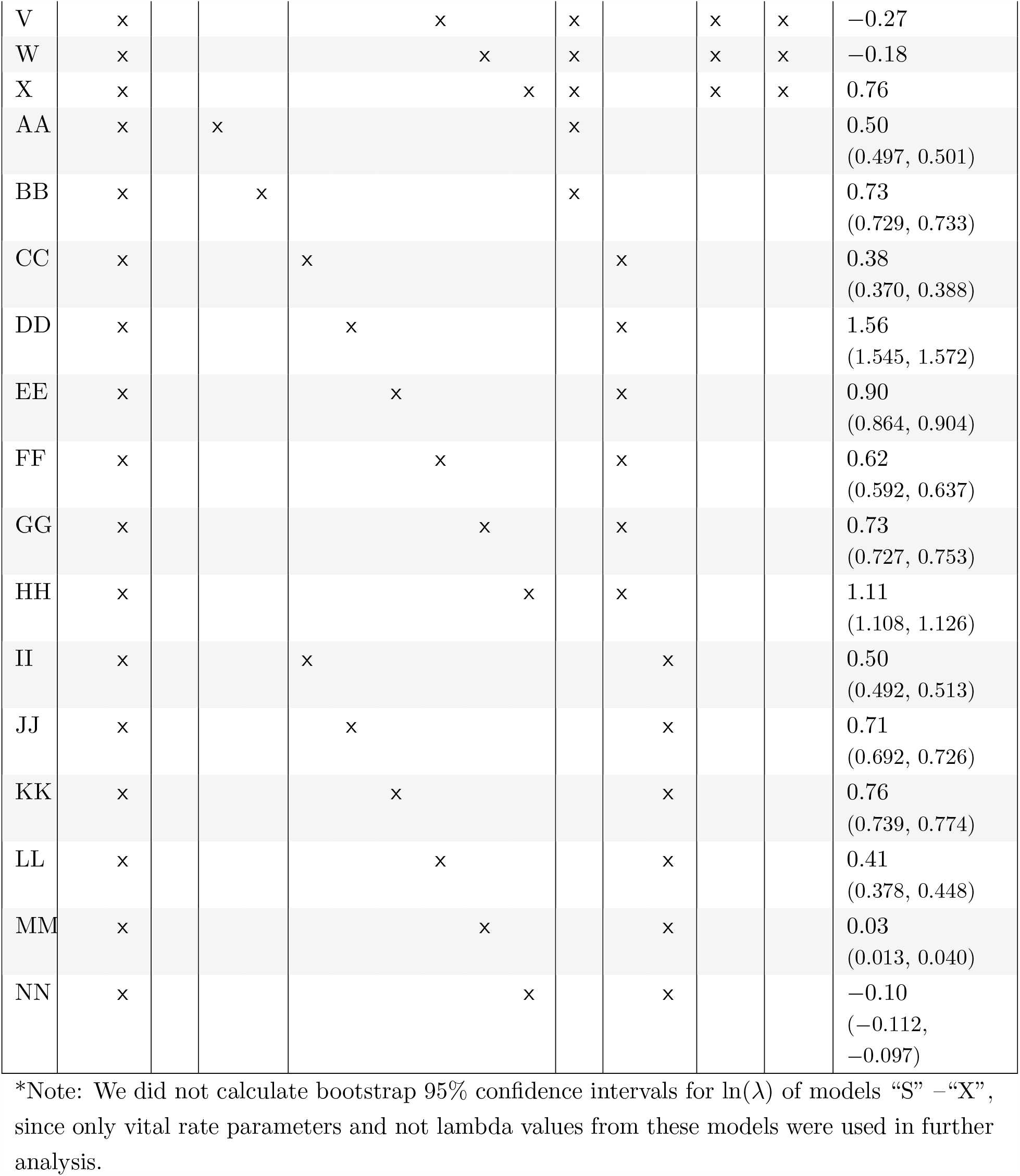
A description of the data used to create each IPM, as well as the covariates included in the vital rate models used in that IPM. ln(*λ*) estimates and 95% bootstrap confidence intervals of ln(*λ*) are also shown for each IPM.

In equations for both types of IPMs, *z* is the distribution of plant longest leaf size (measured as longest leaf length) in the current year (“*size*_*t*_”), *z*′ is the distribution of plant longest leaf size in the next year (“*size*_*t*+1_”), and *U* and *L* are the upper and lower limits of leaf size. *G*(*z*′, *z*) is the vital rate function describing *size*_*t*+1_ as a function of *size*_*t*_. The vital rate functions *s*(*z*), *Pb*(*z*), and *b*(*z*) describe the relationship between *size*_*t*_ and survival probability of non-flowering plants, flowering probability, and seed production of flowering plants, respectively. *c*_*o*_(*z*′) is the distribution of aboveground recruit *size*_*t*+1_. *goCont, outSB, goSB*, and *staySB* are discrete parameters that determine seed bank dynamics. *goCont* is the probability that a seed produced in year *t* germinates as a seedling in year *t* + 1, *outSB* is the probability that a seed from the seed bank in year *t* germinates as a seedling in year *t* + 1, *goSB* is the probability that a seed produced in year *t* goes into the seed bank in year *t* + 1, and *staySB* is the probability of a seed from the seed bank in year *t* persisting in the seed bank in year *t* + 1 (Paniw et al., 2017) (Table 1). *pEstab* is the probability that a seed produced in year *t* establishes as a seedling in year *t* + 1.

We used these vital rate functions and discrete parameters to construct discretized IPM kernels, which were numerically implemented using the “midpoint rule” method (Easterling et al., 2000) with 500 bins, an upper size limit corresponding to 120% of the maximum observed longest leaf size and a lower size limit corresponding to 80% of the minimum simulated seedling size of 0.1 cm. We then used eigen analysis of these kernels to estimate the log-transformed asymptotic population growth rate (ln(*λ*)), damping ratio, stable size distribution, and reproductive value (Caswell, 2001; Ellner, Childs, & Rees, 2016). We used 1000 iterations of bootstrap resampling to estimate 95% bootstrap confidence intervals (95% CIs) for each continuous vital rate parameter included in each IPM, as well as each estimate of ln(*λ*) (Merow et al., 2014; Fieberg, Vitense, & Johnson, 2020). We were unable to estimate CIs for discrete seed bank parameters because they were drawn from a previous publication. We used perturbation analysis to determine the sensitivity and elasticity of ln(*λ*) to changes in germination rate, viability rate, seed survival rate and each continuous vital rate model (Morris & Doak, 2002). Finally, to determine whether including a discrete seed bank state significantly altered our population model, we compared the asymptotic ln(*λ*) and associated 95% CI between IPM “A” and IPM “B.”

### Objective 2: Evaluating Persistence Mechanisms

To evaluate whether any of the demographic mechanisms of rare species persistence outlined in Fig. 1 act in populations of *O. coloradensis*, we fit a series of IPMs that each used different subsets of data as well as additional covariates in vital rate functions to account for density dependence and environmental variation (Table 2: IPMs “C” – “NN”). These IPMs all had a mathematical form equivalent to that of IPM “B” described above, with a discrete seed bank state, and a continuous, size-based stage for above-ground individuals (Eqns. 2 & 3). We then used each of these IPMs, as well as the vital rate functions used to construct them, to evaluate a different persistence mechanism. Details of this process for each persistence mechanism are provided below.

#### Negative Density Dependence

In order to determine the importance of density dependence in *O. coloradensis* subpopulations, we used IPMs and vital rate functions that were fit uniquely for each subpopulation using data from both transitions. However, one set of IPMs included population size in the current year in vital rate models, while another set of IPMs did not (density-independent IPMs: “C”-”H” in Table 2; density-dependent IPMs: “I”-”N”). We used AIC to identify significant differences between vital rate models with and without density dependence terms. We also used results from subpopulation-level IPMs (Table 2: IPMs “CC”-”NN”) for each transition to identify relationships between subpopulation size in *year*_*t*_ and ln(*λ*) (as in Fig. 1), as well as subpopulation size in *year*_*t*_ and the ratio of population size in *year*_*t*+1_ and subpopulation size in *year*_*t*_. In addition to population size information and ln(*λ*) values from our IPMs, we also used population sizes and ln(*λ*) values from a previously-published demographic study of *O. coloradensis* at the three FEWAFB subpopulations that we also monitored (Floyd & Ranker, 1998). A negative relationship between population size in *year*_*t*_ and either ln(*λ*) or the ratio of population size in year *t + 1* to population size in *year*_*t*_ would provide evidence for negative density dependence. Additionally, significant differences between models with and without population size predictor terms would constitute evidence for density dependence.

#### Demographic Compensation

To test for demographic compensation, we calculated the correlation between environmental covariate coefficients in different vital rate models. We used vital rate models that were fit using data from each subpopulation and both transitions, and that included covariates for density dependence and as well as environmental covariates that improved model fit (vital rate models from IPMs “S”-”X” in Table 2). We tested the significance of negative correlations between environmental covariate coefficients using a randomization procedure similar to that used by Villellas et al. (2015), where we randomly assigned an environmental covariate coefficient drawn from the observed distribution of values for that coefficient to each vital rate function, calculated a correlation matrix between those coefficients in each vital rate function, and counted the number of negative correlations in that matrix. We repeated this procedure 10,000 times to generate a null distribution of the expected number of negative correlations between environmental coefficients. We compared the observed number of negative correlations to these expected distributions to determine statistical significance. We could not test for demographic compensation in discrete seed bank vital rate parameters because we did not know how they varied according to environmental conditions. A negative correlation between coefficients for the same covariate in different vital rate models would indicate that demographic compensation was taking place (Villellas et al., 2015; Dibner et al., 2019)(Fig. 1). For example, if soil moisture had a positive effect on growth but a negative effect on survival, this would be evidence for demographic compensation.

#### Vital Rate Buffering

We tested for the presence of vital rate buffering in *O. coloradensis* populations by comparing the variability of vital rates to their importance. We used an approach that scales both the standard deviation (variability metric) and sensitivity (importance metric) of vital rates, allowing for a fair comparison of variability and importance across vital rates with fundamentally different relationships between their mean and variance (McDonald et al., 2017). Vital rates that are probabilities are constrained between zero and one and thus typically have small variance as the mean approaches these limits, while other vital rates are only constrained by zero and thus typically have variances that increase as the mean increases (Gaillard & Yoccoz, 2003). To enable a fair comparison between these different categories of vital rates, we calculated their importance and variability metrics in different ways. Importance of probability vital rates corresponded to the logit variance stabilized sensitivity, and variability corresponded to the standard deviation of the logit transformed vital rate values. Importance of non-probability vital rates corresponded to the log-scaled sensitivity (or elasticity), and variability was corresponded to standard deviation of the log-transformed vital rate values (McDonald et al., 2017; Morris & Doak, 2002; Link, William A, Paul F Doherty, Fr., 2002).

We used IPM “B” to calculate elasticity or logit VSS values for each discrete and continuous vital rate. We calculated the scaled standard deviation for each continuous vital rate function using the vital rates that were fit uniquely for each subpopulation and each transition (Table 2: IPMs “CC”-”NN”). Because we did not have site-level information about discrete seed bank vital rates, we simulated both the maximum and minimum possible standard deviations for each discrete vital rate. We then proceeded with two comparisons of vital rate variability and importance, one using the maximum possible discrete vital rate standard deviation, and another using the minimum. In order to determine the correlation between a single importance/variability value pair for discrete vital rates and a string of value pairs for continuous vital rate functions, we calculated mean importance and variability values for each continuous vital rate function. A significant negative correlation between the mean or absolute scaled importance (logit VSS or elasticity) and mean or absolute variability (standard deviation of logit or log-transformed vital rates) across all vital rates would constitute support for the presence of vital rate buffering in this species (Fig. 1).

#### Asynchronous Responses and Source-Sink Dynamics

To determine whether *O. coloradensis* subpopulations showed asynchronous responses to environmental variation, we made a correlation matrix to determine how change in ln(*λ*) from year to year was correlated across each subpopulation, using values of ln(*λ*) derived from IPMs for each subpopulation (Table 2: IPMs “C”-”H”). We used the “mantel()” function from the “vegan” R package to perform a Mantel test, which determined if the Spearman correlation of ln(*λ*) across subpopulations was significantly related to the Euclidean distance between each subpopulation (Oksanen et al., 2020). A negative relationship between the distance between subpopulations and degree of correlation of ln(*λ*) would constitute evidence for spatial asynchrony between subpopulations (Fig. 1).

Because we did not have information about gene flow between subpopulations of *O. coloradensis* via pollination or seed dispersal, it was impossible to directly measure whether fine-scale source-sink dynamics were acting in these populations. However, because variation in ln(*λ*) across space is a prerequisite for source-sink dynamics, the tests for spatial asynchrony in subpopulations can also provide evidence for the existence of source-sink dynamics. Again, this would be a negative relationship of distance between subpopulations and correlation of subpopulation ln(*λ*) (Fig. 1).

## Results

### Vital Rate Models

Larger non-reproductive plants were more likely to survive than smaller plants (Fig. 3A). Plants below ∼ 7.5 cm were likely to be larger, while plants larger than ∼ 7.5 cm were likely to be smaller the following year (Fig. 3 B). Flowering probability was best approximated as a quadratic polynomial, where flowering probability peaked at 12 cm leaf length, and plants with the largest leaves exhibited low flowering probability (Fig. 3 C). The number of seeds that a reproductive plant produced increased sharply with leaf size (Fig. 3 D). The inclusion of additional covariates did not alter the overall shape or sign of the relationships between leaf size and vital rates, so models shown in Figure 3 did not include any additional covariates beyond leaf size.

### Objective 1: Importance of the Seed bank Stage

#### Integral Projection Models

We found that including a discrete seed bank stage in IPMs for *O. coloradensis* significantly increased ln(*λ*). The continuous state-only IPM (Table 2: IPM “A”) predicted an asymptotic ln(*λ*) of 0.27 for all populations (95% CI: 0.269 - 0.271), while the continuous + discrete state IPM (Table 2: IPM “B”) predicted an asymptotic ln(*λ*) of 0.65 (populations (95% CI: 0.648 - 0.650). All subsequent IPM results refer to models that included a discrete seed bank state.

The simplest two-state IPMs that excluded density dependence and environmental variation indicated that both the Soapstone and FEWAFB populations had positive ln(*λ*) values (Table 2: Soapstone - IPM “AA”, ln(*λ*) = 0.50; FEWAFB – IPM “BB”, ln(*λ*) = 0.73). The Diamond Creek subpopulation at FEWAFB had the highest ln(*λ*) from 2018 to 2020 (Table 2: IPM “D”, ln(*λ*) = 1.13), while the HQ3 subpopulation at Soapstone had the lowest growth rate (Table 2: IPM “G”, ln(*λ*) = 0.395). Almost all additional IPMs identified a positive ln(*λ*) (Table 2).

A density-independent, discretized IPM kernel (made using IPM “B” in Table 2) calculated transition probabilities within and between the discrete and continuous stages of the *O. coloradensis* life cycle when all populations and transitions were considered together (Fig. 4 A). Relative to other demographic transitions, there was a high probability that seeds stayed in the seed bank, and a high probability that seeds from medium-sized adult plants go into the seed bank in the next year. Population *λ* was most sensitive to the rates at which seeds were produced by adult plants and stayed in the seed bank(Fig. 4 C).

**Figure 3.**
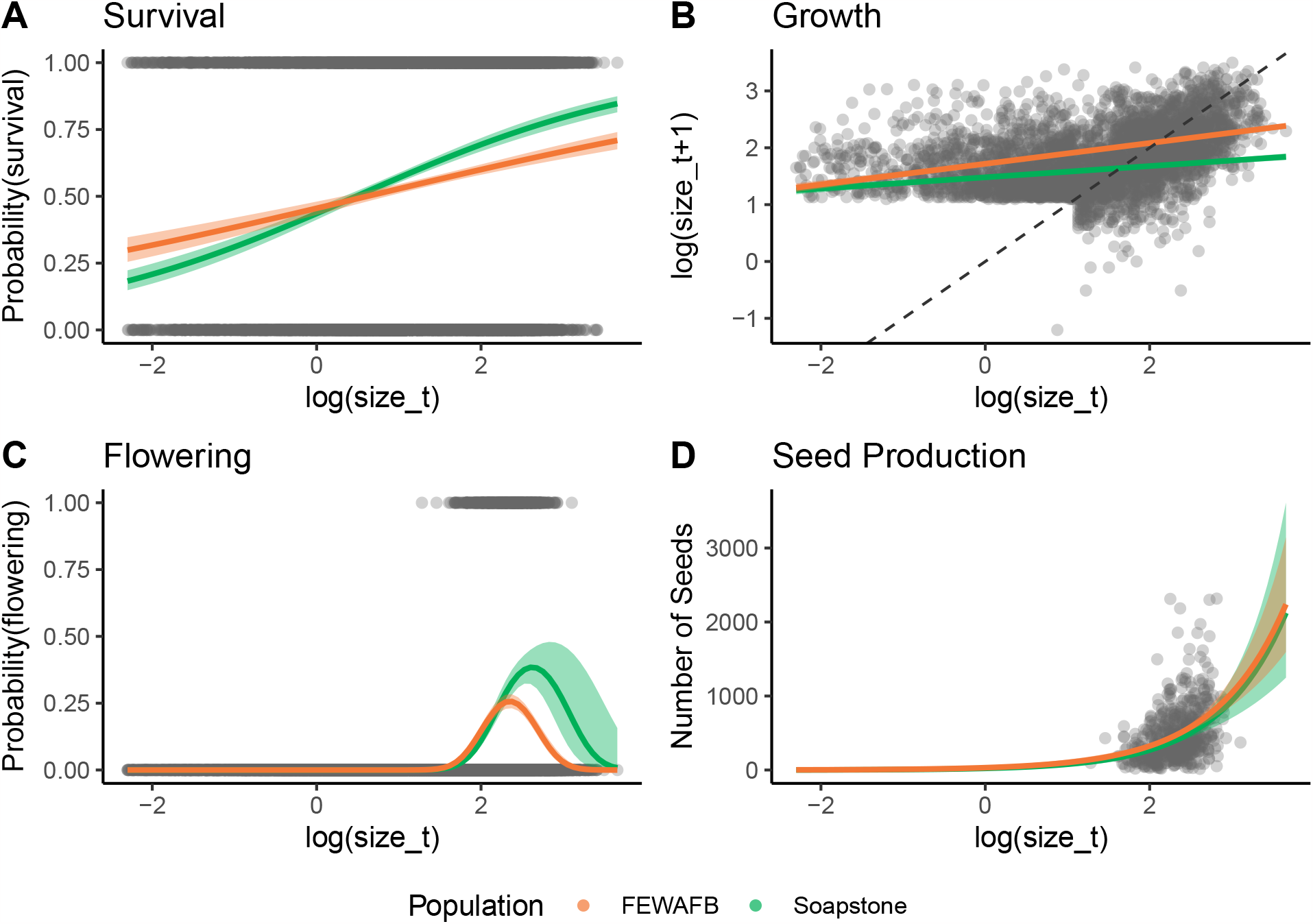
The effect of current year leaf size (ln(size_*t*_)) on vital rates in monitored *O. coloradensis* populations. Data for from all sites and all transitions are shown. Lines indicate vital rate functions for each population, fit using the “global” model structure described in the Methods section. Bands around each line show 95% confidence intervals. The dashed line in panel **B** shows a 1:1 line. The sharp cut-off in ln(size_*t*+1_) in panel **B** is due to the fact that two-year-old plants could not be seedlings, which were classified as any plant less than 3 cm in size.

### Objective 2: Evaluating Persistence Mechanisms

#### Negative Density Dependence

We found evidence that negative density-dependence occurred in subpopulations of *O. coloradensis*. AIC comparison of continuous vital rate models indicated that density-dependent models were better predictors of the majority of vital rates than density-independent models in most subpopulations (Table 3). Models that included population size in the previous year as a covariate were better predictors of growth in five of six subpopulations. Density dependent models were better predictors of survival and seed production in four out of six subpopulations, and density dependent models of flowering were better in one subpopulation. Recruit size distribution was not affected by density dependence. The vital rate models for the Meadow population at Soapstone were least affected by density dependence. Although density dependence was important for *O. coloradensis* in many situations, it appeared only to be acting to decrease *λ* at high density (as in the highly dense Diamond Creek or HQ5 subpopulations), but not clearly increasing lambda at low density (as in the sparsely populated Meadow subpopulation). We also found that, within a subpopulation, ln(*λ*) was generally higher when subpopulation size was smaller(Fig. 5 A). Similarly, there was a negative relationship within each subpopulation between subpopulation size in *year*_*t*_ and the ratio of subpopulation size in *year*_*t*+1_ to subpopulation size in *year*_*t*_ (Fig. 5 B and D); Interestingly, these negative relationships are generally pronounced at the subpopulation level, but are weak when examining data across all subpopulations.

**Figure 4.**
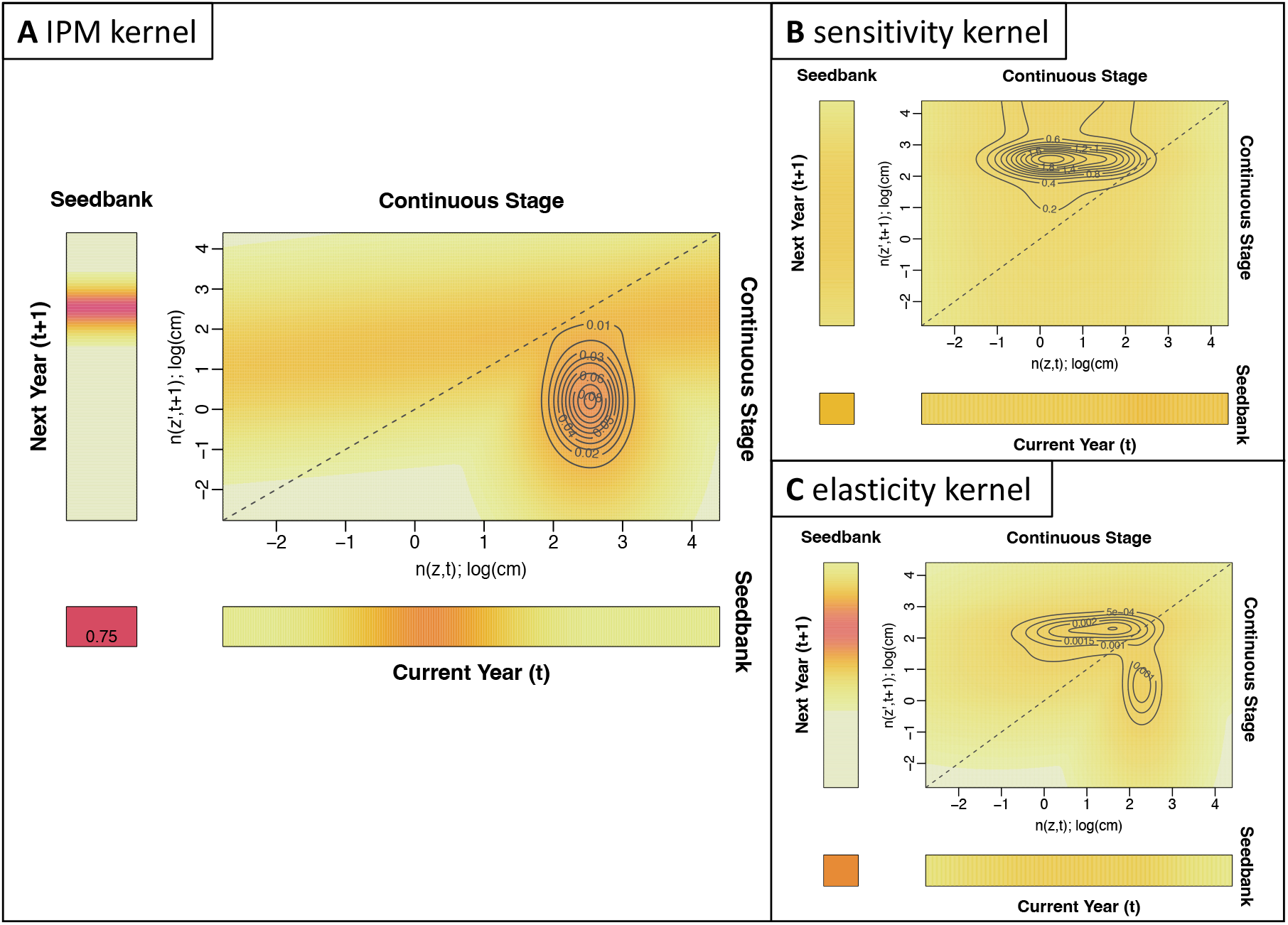
Visualizations of the *O. coloradensis* IPM kernels. (**A**) The IPM kernel for *O. coloradensis*. This kernel shows a density-independent IPM constructed using all data from all transitions (IPM “B”). (**B**) Sensitivity of the IPM kernel. (**C**) Elasticity of the IPM kernel. In all panels, color indicates probability, with darker colors corresponding to higher probability, and lighter colors corresponding to lower probability. The dashed line shows a 1:1 line.

#### Demographic Compensation

We did not find evidence of demographic compensation in *O. coloradensis* populations. While there were negative correlations between the effect of mean growing season temperature on vital rates for five combinations of vital rates, none of these correlations were significant (Table 4). The only significant correlation, between temperature coefficients in growth and survival models, was positive. Ten thousand correlations of randomly assigned coefficients found that the number of negative correlations in a matrix can be described by a normal distribution with a mean of 4.97 and a standard deviation of 1.60. Using this distribution as a null model, there was a 50.7% probability of observing five negative correlations. Although there is no significant evidence for demographic compensation, it is notable that the effect of mean growing season temperature on distribution of recruit size was negatively correlated with the effect of growing season temperature on all other vital rates. We were only able to compare coefficients across vital rate models for mean growing season temperature, because including precipitation and mean winter temperature as covariates resulted in overfitting in some cases.

#### Vital Rate Buffering

We did not find evidence of vital rate buffering in the *O. coloradensis* populations we observed. Vital rate importance (either logistic VSS or elasticity) and variability (corrected SD) were not significantly negatively correlated, regardless of the simulated standard deviation for discrete vital rates we used(Fig. 6; correlation with minimum discrete vital rate SD (**A**): *r* = 0.43, *P* = 0.25; correlation with maximum discrete vital rate SD (**B**): *r* = -0.07, *P* = 0.85). As a vital rate became more important for determining ln(*λ*), it did not become significantly less variable, as it would if vital rate buffering was occurring (Fig. 1).

**Table 3:** Comparison of vital rate models with and without density dependence. Values show the ∆AIC, or difference between the AIC of density independent (DI) and density dependent (DD) models. Bold text indicates that the |∆|AIC value is *>* 3, which means that including a term for density dependence substantially changed that vital rate model. A positive |∆|AIC indicates that including density dependence improved the model, while a negative value indicates that including density dependence made model fit worse. AIC values for DI and DD values can be found in Table S3.

| Vital Rate Model | Subpopulation Data |  |  |  |  |  |
| --- | --- | --- | --- | --- | --- | --- |
|  | Crow Creek | Diamond Creek | Unnamed Creek | HQ5 | HQ3 | Meadow |
| Survival | **18.74** | **107.28** | -0.41 | **320.33** | **78.82** | -0.70 |
| Growth | **3.73** | **22.15** | **30.811** | **458.15** | **30.45** | 2.66 |
| Flowering | -1.63 | -0.44 | 0.45 | **54.52** | -1.76 | -1.72 |
| Seed production | **6.41** | **14.02** | -1.29 | **3.71** | **4.12** | -1.35 |
| Recruit size | -1.93 | 1.61 | -1.93 | -1.97 | -1.23 | -1.99 |

**Figure 5.**
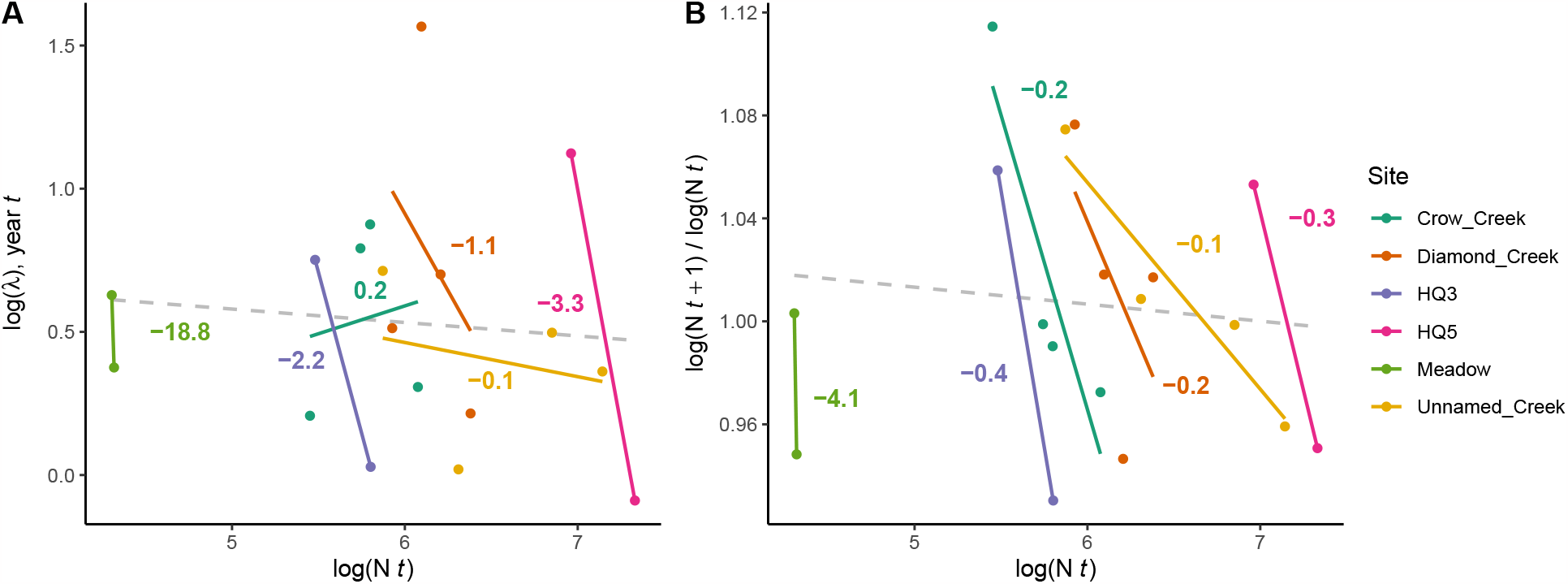
(**A**) Within the same subpopulation, ln(*λ*) calculated from IPMs decreases as population size increases. (**B**) Within each subpopulation, ln(*λ*) calculated by change in population size from *year*_*t*_ to *year*_*t*+1_ also decreases as population size increases. In **A** and **B**, each point represents values calculated from data from one transition in one subpopulation. Solid lines show linear regressions of the relationships between ln(N_*t*_) and the respective response variable in each subpopulation. Numbers adjacent to these solid lines show the slope of each relationship, and are color-coded by subpopulation. Dashed grey lines show linear regressions of the relationships between ln(N_*t*_) and the respective response variable across all subpopulations.

**Figure 6.**
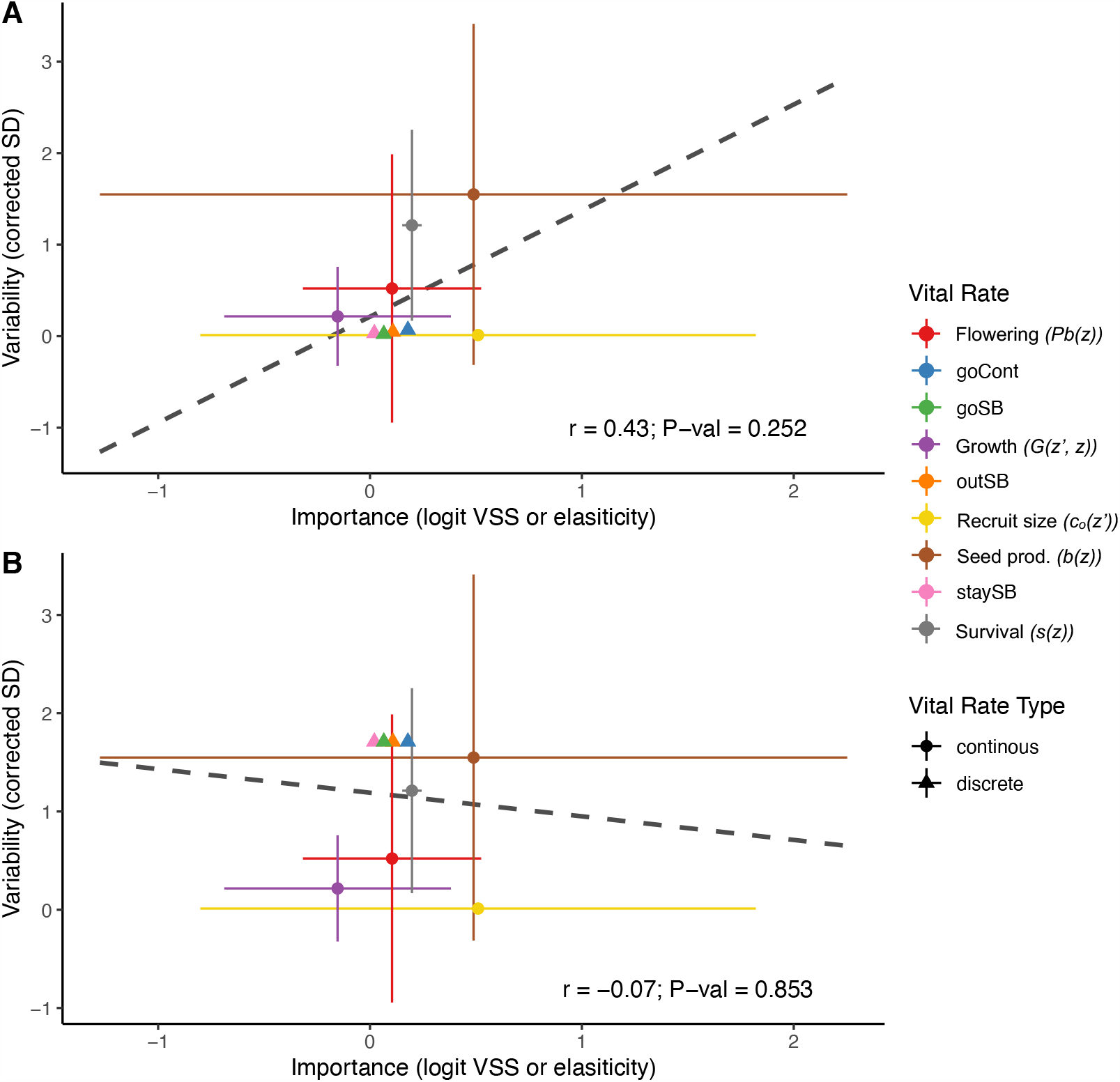
The relationship between variability of each vital rate (measured by corrected standard deviation) and its importance (measured by logit VSS or elasticity) does not show support for vital rate buffering. (**A**) With the minimum posible simulated discrete vital rate variability, there is a positive but insignificant correlation between vital rate variability and importance (*r* = 0.43, *P* = 0.25). (**B**) Using the maximum possible simulated discrete vital rate variability, there is a negative but insignificant correlation between vital rate variability and importance (*r* = -0.07, *P* = 0.85). Triangles indicate importance and variability for discrete vital rate parameters, while circles indicates the means of importance and variability across an entire continuous vital rate function. Error bars around continuous vital rate means span the the 5^*th*^ to 95^*th*^ percentiles of either importance or variability values calculated for an entire continuous vital rate function. Dashed lines show the correlation between (mean) variability and (mean) importance across all vital rates.

#### Asynchronous Responses and Source-Sink Dynamics

We did not find evidence of asynchronous responses to environmental variation in *O. coloradensis* populations. There was not a significant relationship between the Spearman correlation of ln(*λ*) between subpopulations and their spatial proximity (Mantel statistic = 0.396, *P* = 0.06). We also performed Mantel tests using ln(*λ*) correlation and distance matrices calculated uniquely for each population, and did not find evidence for asynchronous responses (Soapstone: Mantel statistic = -0.659, *P* = 0.83; FEWAFB: Mantel statistic = 0.798, *P* = 0.33). We did, however, find a positive relationship between correlation of ln(*λ*) and distance between subpopulations at Soapstone, and a negative relationship between sub-populations at FEWAFB. Collectively, these results fail to provide support for both asynchronous responses and fine-scale source-sink dynamics in these *O. coloradensis* populations.

**Table 4:**
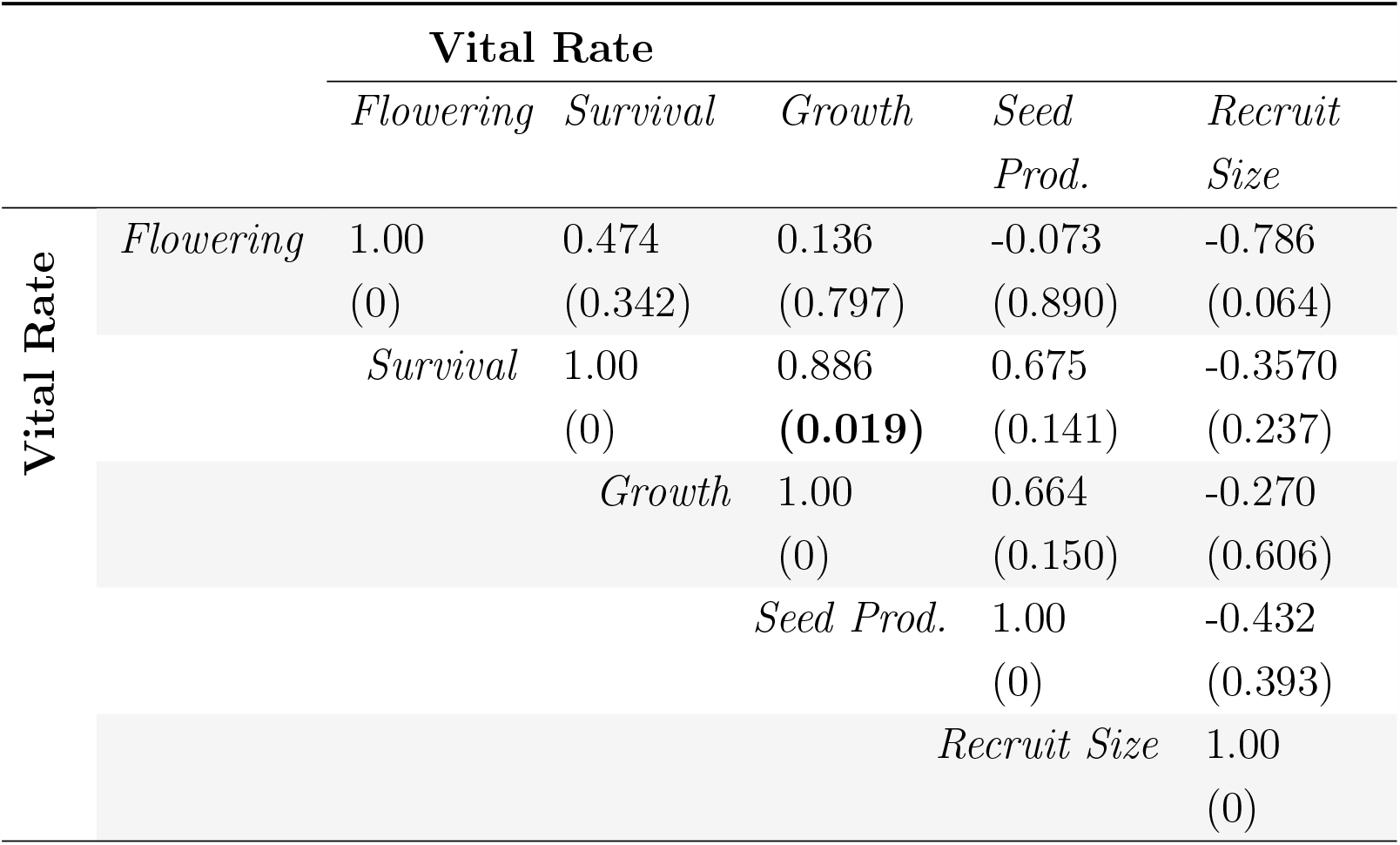
Pearson correlations between mean growing season temperature coefficients in each continuous vital rate function. Below each correlation value is the *P* value for that correlation. Bold text indicates a significant correlation.

|  |  | Vital Rate |  |  |  |  |
| --- | --- | --- | --- | --- | --- | --- |
|  |  | *Flowering* | *Survival* | *Growth* | *Seed Prod.* | *Recruit Size* |
| Vital Rate | *Flowering* | 1.00<br>(0) | 0.474<br>(0.342) | 0.136<br>(0.797) | -0.073<br>(0.890) | -0.786<br>(0.064) |
|  | *Survival* |  | 1.00<br>(0) | 0.886<br>**(0.019)** | 0.675<br>(0.141) | -0.3570<br>(0.237) |
|  | *Growth* |  |  | 1.00<br>(0) | 0.664<br>(0.150) | -0.270<br>(0.606) |
|  | *Seed Prod.* |  |  |  | 1.00<br>(0) | -0.432<br>(0.393) |
|  | *Recruit Size* |  |  |  |  | 1.00<br>(0) |

## Discussion

Our demographic analysis of the two largest known populations of the globally rare *Oenothera coloradensis* evaluated the importance of seed banks to population dynamics and the demographic mechanisms that allow this rare species to persist. We found that including information about cryptic life stages alters the outcomes of the population model (Paniw et al., 2017; Nguyen et al., 2019), and that *O. coloradensis* populations show signs of negative density-dependence at the subpopulation scale (Fig. 5; Table 3). However, these populations do not show substantial evidence of demographic compensation, vital rate buffering, spatial asynchrony, or fine-scale source-sink dynamics. This may indicate that while these mechanisms may be important for the persistence of many small populations of rare plants, they are not strictly necessary in all cases.

Including a discrete seed bank state in an IPM increased the asymptotic population growth rate (ln(*λ*)) compared to an IPM with only a continuous, size-based state, although both growth rates were still positive (Table 2: with seed bank: IPM “B”, ln(*λ*) = 0.65; without seed bank: IPM “A”, ln(*λ*) = 0.27). The importance of including the seed bank in the model was consistent with our expectations, and also aligns with the conventional notion that seed banks can act as buffers against stochastic causes of population decline. The discrete rates for the probability of persisting and transitioning out of the seed bank have high elasticity in the IPMs in which they are included, but not the highest elasticity of any vital rate(Fig. 4 C). The rate at which seeds produced by adult plants in *year*_*t*_ go into the seed bank in *year*_*t*+1_ is the vital rate function with highest elasticity. Previous matrix population models of *O. coloradensis* without a seed bank state that were constructed in the 1990s identified the emergence rate of new seedlings as the vital rate most important for determining ln(*λ*) (Floyd & Ranker, 1998). Our finding that seed bank state transitions are important for this species aligns with this previous result, since rate of seedling emergence is the above-ground plant vital rate that is closest to the seed bank in this plant’s life cycle. An important caveat to our comparison of models with and without seed bank stages is the fact that the seed bank vital rate parameters we used were inferred from laboratory tests of germination and viability rates, which may be imperfect representations of *in-situ* rates of viability and germination. The annual rate of seed death (10%) was inferred from an *in-situ* study, but is likely imprecise because of low sample size. Regardless of these potential sources of error, our results reinforce the fact that the seed bank can be an important element of a perennial plant’s life cycle, and if possible, should be modeled explicitly based on *in-situ* estimates of the probability of seeds going into, persisting in, and emerging from the seed bank.

We found evidence that, of the five proposed demographic mechanisms of small population persistence, negative density dependence was the only one acting in these *O. coloradensis* populations. Including population size in the previous year as a covariate in vital rate models typically improved model fit, suggesting that density dependence is an important driver of growth, survival, and reproduction (Table 3). Within a single subpopulation, ln(*λ*) and the ratio of population size in *year*_*t*+1_ to *year*_*t*_ was generally higher when population size in *year*_*t*_ was smaller (Fig. 5), which indicates that negative density dependence prevents subpopulations from crashing when their population size is very small. However, this pattern of higher growth rate at low population sizes did not exist when considering all subpopulations together (Fig. 5). This could indicate that each subpopulation is close to its carrying capacity for *O. coloradensis*. This may indicate that the number of individuals is close to carrying capacity in each subpopulation, and that growth rate increases when the population size in a given subpopulation is small in comparison to its subpopulation-specific carrying capacity. *O. coloradensis* vital rates had correlated responses to variation in the abiotic environment (Table 4), which is the inverse of what is expected if demographic buffering is taking place. It is possible that a signal of demographic buffering would appear if we considered different abiotic variables such as disturbance frequency, or had more data. Vital rate buffering also was not identified, either with the minimum or maximum possible simulated discrete vital rate variability (Fig. 6). Vital rates with higher variability (higher SD) did not have a significantly higher or lower importance for determining ln(*λ*) in comparison to less variable vital rates. This indicates that vital rate buffering is not stabilizing ln(*λ*) after abiotic or demographic perturbation. The evidence for spatial asynchrony and fine-scale source-sink dynamics was also not strong. Mantel tests did not identify a significant relationship between the correlation of ln(*λ*) between subpopulations and their spatial proximity, but did identify non-significant relationships between ln(*λ*) correlation and proximity. However, this relationship was positive in Soapstone prairie subpopulations and negative in FEWAFB subpopulations, which provides inconsistent support for these mechanisms.

It is somewhat surprising that negative density dependence is the only mechanism of small population persistence that has significant support in *O. coloradensis* populations, since multiple mechanisms have been identified in at least one other rare species (Dibner et al., 2019). It is possible that support for one or more of these persistence mechanisms could emerge if more information about abiotic variation across space and time and data from more annual transitions was available for analysis. One potential explanation is that, while this species is a globally rare endemic with isolated subpopulations, it often grows at high local density. This strategy, which Rabinowitz describes as “locally abundant in a specific habitat but restricted geographically,” may allow *O. coloradensis* to bypass the problems that small populations typically face, such as genetic and demographic bottlenecks that make them susceptible to stochastic environmental variation (Rabinowitz, 1981). It has also been shown that rare species are more likely than common species to benefit from facilitative interspecific interactions (Calatayud et al., 2020). *O. coloradensis* may participate in facilitative interactions with other species that increase its probability of persistence, although determining this will require further, community-level analysis. Our results imply that not all rare species can be treated equally. While demographic strategies that help maintain persistence may be effective for some species, other species may employ different strategies. This further emphasizes the importance of carefully considering the specific population and its community dynamics when managing and conserving rare species.

Our analysis of the population dynamics of *Oenothera coloradensis* at two distinct locations shows that this species has a life cycle that is strongly driven by introduction and persistence of seeds into a seed bank. More broadly, we show that this rare endemic species shows signs of negative density dependence. Populations of *O. coloradensis* may additionally be maintained via high local abundances that allow them to escape the challenges of small population size that rare species often face (Rabinowitz, 1981). These findings reinforce the importance of careful evaluation of the unique population dynamics of rare species to inform successful conservation and management.

## Acknowledgements

Funding for this project was provided by the Wyoming Native Plant Society Markow Grant, and the U.S. Fish and Wildlife Service (USFWS) through a grant to the Wyoming Natural Diversity Database and the University of Wyoming Botany Department (grant #14AC00827). Collection of seeds for germination experiments was conducted under USFWS Permit #TE085324-2. Data at Soapstone Prairie were collected under permits #1082-2018 (2018), #1166-2019 (2019), and #4919647-26 (2020). Data collection at F.E. Warren Air Force Base was conducted with permission and assistance from Alex Schubert, USFWS Fish and Wildlife Biologist.

