## Supplemental Information for "Negative density dependence promotes persistence of a globally rare yet locally abundant plant species (*Oenothera coloradensis*)"

### Supplementary Materials

#### Species Information

*Oenothera coloradensis* seeds are contained within small, indehiscent capsules that contain 2-5 seeds each (Burgess, Hild, & Shaw, 2005). A single adult individual can produce >500 capsules. This species does not reproduce vegetatively, although seeds typically germinate near the base of the parent plant, which often results in dense patches of vegetative individuals (Heidel, Tuthill, & Wallace, 2021). *O. coloradensis* is pollinated primarily by hawkmoths (Krakos, pers. comm. to B. Heidel, 2013). Seed dispersers are unknown (Floyd & Ranker, 1998; Heidel et al., 2021). Previous work established that *O. coloradensis* population growth rate is particularly impacted by recruitment of seedlings (Floyd & Ranker, 1998). Recruitment increases when non-*O. coloradensis* community biomass is removed, indicating that surrounding grasses and forbs outcompete or shade-out seedlings (Munk, Hild, & Whitson, 2002).

*O. coloradensis* commonly co-occurs with *Agrostis stolonifera*, *Pascopyrum smithii*, *Poa pratensis*, *Glycyrrhiza lepidota*, *Iris missouriensis*, *Cirsium flodmanii*, and *Grindelia squarrosa* (Endangered and Threatened Wildlife and Plants, 2000; Munk et al., 2002). Encroachment of woody shrubs such as *Salix exigua* has been correlated with declining numbers in some populations (Heidel et al., 2021).

The Wyoming Natural Diversity Database (WYNDD) began a base-wide census of reproductive individuals in the FEWAFB population in 1986, and has repeated this census annually since 1988 (Heidel et al., 2021). The first estimate of species size after its full geographic range was identified occurred in 1998, when it was approximated that the entire species consisted of 47,300 to 50,300 reproductive individuals (Fertig, 2000).

### Seed Production Estimation

It was not possible to measure seed production exactly because O. coloradensis seeds are contained in indehiscent capsules. Additionally, buds on the same individual flower and set seed with a time lag of up to several weeks, so mature seed capsules often exist at the tip of a stem while un-opened buds lower down on that same stem have not yet flowered. This lag makes it difficult to count the total number of capsules produced by an individual. However, seed capsules leave a noticeable scar on the stem, so we used the number of seed capsule scars on reproductive stems as an estimate of capsule production. Counting scars is extremely time-intensive since a single plant can produce several hundred capsules, so we used Poisson generalized linear regression to estimate the relationship between the length of stem bearing capsule scars and the number of capsules produced by that stem. A Poisson regression model fit to stem measurements and capsule counts from 106 individuals in 2018 indicated that the number of capsules produced by an individual (C) can be predicted by $e^{(1.843+0.119\text{x}S))}$, where S is the stem length in cm (pseudo $R^2 = 0.42$, P $< 0.01$, Residual deviance = 186.98, df = 104) (Fig. S2). We used this relationship to estimate capsule production for each reproductive individual. Previous work indicated that each capsule contained an average of 4 seeds, so we multiplied the estimated number of capsules produced by an adult plant by 4 to estimate seed production (Burgess et al., 2005).

### Discrete Vital Rate Parameters

Previously-published data from a greenhouse experiment using *O. coloradensis* seed capsules collected from the FEWAFB populations determined that viable seeds had an average germination rate of 20.3% after cold-stratification, and did not identify a consistent decline in germination rate over five years (Burgess et al., 2005). This study also found that 58.5% of seeds produced were viable. We conducted an additional seed study to determine if overwintering in natural conditions lead to a lower germination rate than was identified in the previous greenhouse study. We buried 60 field-collected seed capsules in mesh bags at 6 locations near our demographic study plots at FEWAFB, and then recovered the seed bags after one winter. An average of 10% of seed capsules were not recoverable, likely because they were non-viable and withered away or were eaten. We planted the recovered capsules in standard greenhouse conditions, and found a mean germination rate of 6.8%. This germination rate was much lower than that identified by Burgess et al., however our seed study had a much smaller sample size, reducing the reliability of our result (Burgess et al., 2005). However, it is still likely that true germination rates are much lower than those identified in greenhouse conditions, so we reduced the germination rate identified by Burgess et al. by 20%.

### Seedling Data

Although seedlings (above-ground plants $< 3$ cm in leaf length) were only tallied in each plot quadrant and year instead of tagged and measured, we incorporated them into the dataset for continuous, above-ground plants by assigning them a random size drawn from a continuous, uniform probability distribution (seedling size $\sim U(0.1, 3)$). Each new recruit to the $> 3$ cm stage in year $t + 1$ was randomly assigned to a seedling within the same plot quadrant in year $t$. Seedlings in year $t$ that were assigned a recruit in year $t + 1$ survived, while those without an assigned recruit died. Incorporating seedlings into the continuous dataset in this fashion allowed us to create IPMs using only one discrete stage.

Table S1: Permanent Plot Locations and subpopulation-level sample sizes for each year and individual type (seedling vs. non-seedling). GPS coordinates listed in decimal degrees, map datum and spheroid: WGS 84.

| | | | | | Sample Size | | | | | |
|---|---|---|---|---|---|---|---|---|---|---|
| | | | | | 2018 | | 2019 | | 2020 | |
| Site | Subpopulation | Plot Name | N Coord. | W Coord. | non-seedling | seedling | non-seedling | seedling | non-seedling | seedling |
| FEWAFB | Unnamed Creek | U3 | 41.13642 | -104.87209 | | | | | | |
| | Unnamed Creek | U4 | 41.13634 | -104.87183 | 740 | 525 | 528 | 417 | 406 | 530 |
| | Unnamed Creek | U6 | 41.13647 | -104.87132 | | | | | | |
| | Diamond Creek | D7 | 41.14340 | -104.88380 | | | | | | |
| | Diamond Creek | D10 | 41.14441 | -104.88303 | 235 | 209 | 347 | 149 | 275 | 81 |
| | Diamond Creek | D11 | 41.14431 | -104.88094 | | | | | | |
| | Crow Creek | C4 | 41.15540 | -104.87497 | | | | | | |
| | Crow Creek | C5 | 41.15477 | -104.87474 | 203 | 127 | 214 | 98 | 150 | 160 |
| | Crow Creek | C8 | 41.15534 | -104.87487 | | | | | | |
| Soapstone | Pasture HQ5 | S1 | 40.99297 | -105.00925 | | | | | | |
| | Pasture HQ5 | S2 | 40.99318 | -105.00935 | 283 | 772 | 714 | 813 | 641 | 423 |
| | Pasture HQ5 | S3 | 40.99342 | -105.00937 | | | | | | |
| | Pasture HQ3 | S4 | 40.98623 | -105.01691 | | | | | | |
| | Pasture HQ3 | S5 | 40.98639 | -105.01671 | 102 | 138 | 158 | 173 | 117 | 104 |
| | Pasture HQ3 | S6 | 40.98650 | -105.01656 | | | | | | |
| | Meadow | S7 | 40.98753 | -105.02148 | | | | | | |
| | Meadow | S8 | 40.98747 | -105.02179 | 44 | 31 | 47 | 28 | 48 | 12 |
| | Meadow | S9 | 40.98724 | -105.02145 | | | | | | |

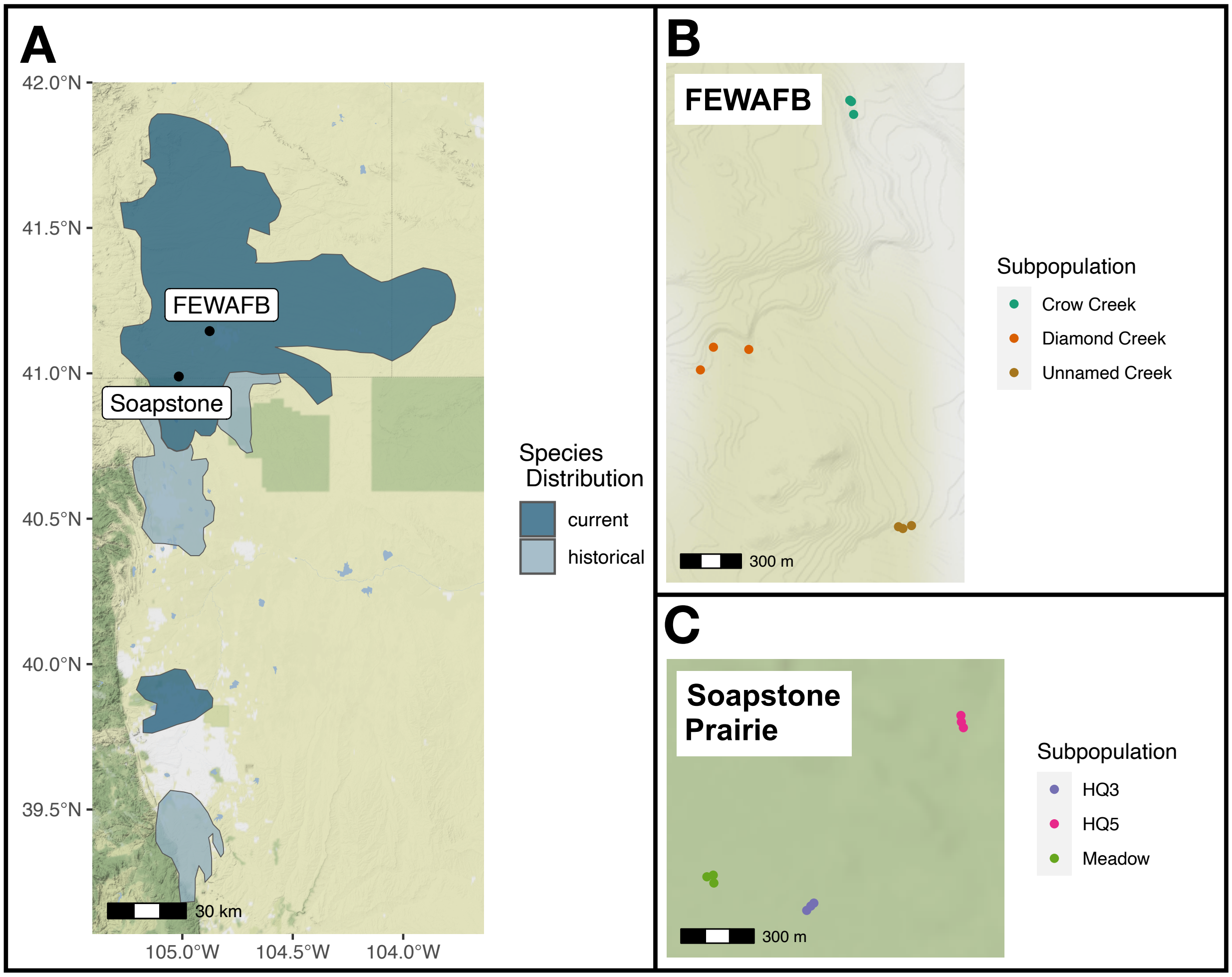


Figure S1: (**A**) The current known distribution of *O. coloradensis*, shown in dark blue, extends into Wyoming, Colorado, and Nebraska. The historical distribution included the current distribution area as well as some additional locations shown in pale blue. Distribution information comes from Everson, 2019. Black dots show the relative locatino of the FEWAFB and Soapstone prairie populations included in this study. Colored dots show the location of plots in each subpopulation at FEWAFB (**B**) and Soapstone Prairie (**C**).

Table S2: Continuous vital rate functions used in each IPM. In the functions below, $size_t$ indicates plant longest leaf length in the previous year, $N_t$ indicates number of individuals in a plot in the previous year, $T_G$ indicates mean temperature in the previous year's growing season, and $T_W$ indicates the mean temperature in winter of the previous year.

| IPM | Vital Rate | Equation |
|---|---|---|
| A & B | Survival | $logit(s(z)) = -0.21 + 0.37(ln(size_t))$ |
| | Flowering | $\text{logit}(Pb(z)) = -34.06 + 27.67(ln(size_t)) - 5.78(ln(size_t)^2)$ |
| | Growth | $G(z', z) = N(\mu_s, \sigma_s)$; $\mu_s = 1.57 + 0.18(ln(size_t))$; $\sigma_s = 0.51$ |

| | | |
|---|---|---|
| | Seed prod. | $exp(b(z)) = 3.35 + 1.19(ln(size_t))$ |
| | Recruit size | $c_o(z') = N(\mu_r, \sigma_r)$; $\mu_r = 0.21$; $\sigma_r = 0.77$ |
| C | Survival | $logit(s(z)) = -0.73 + 0.45(ln(size_t))$ |
| | Flowering | $logit(Pb(z)) = -40.31 + 33.40(ln(size_t)) - 7.09(ln(size_t)^2)$ |
| | Growth | $G(z', z) = N(\mu_s, \sigma_s)$; $\mu_s = 1.64 + 0.11(ln(size_t))$; $\sigma_s = 0.42$ |
| | Seed prod. | $exp(b(z)) = 3.73 + 1.01(ln(size_t))$ |
| | Recruit size | $c_o(z') = N(\mu_r, \sigma_r)$; $\mu_r = 0.19$; $\sigma_r = 0.76$ |
| D | Survival | $logit(s(z)) = 1.01 + -0.01(ln(size_t))$ |
| | Flowering | $logit(Pb(z)) = -44.52 + 36.73(ln(size_t)) - 7.69(ln(size_t)^2)$ |
| | Growth | $G(z', z) = N(\mu_s, \sigma_s)$; $\mu_s = 1.84 + 0.26(ln(size_t))$; $\sigma_s = 0.52$ |
| | Seed prod. | $exp(b(z)) = 3.80 + 1.11(ln(size_t))$ |
| | Recruit size | $c_o(z') = N(\mu_r, \sigma_r)$; $\mu_r = 0.22$; $\sigma_r = 0.78$ |
| E | Survival | $logit(s(z)) = 0.21 + 0.18(ln(size_t))$ |
| | Flowering | $logit(Pb(z)) = -18.73 + 15.84(ln(size_t)) - 3.63(ln(size_t)^2)$ |
| | Growth | $G(z', z) = N(\mu_s, \sigma_s)$; $\mu_s = 1.74 + 0.24(ln(size_t))$; $\sigma_s = 0.49$ |
| | Seed prod. | $exp(b(z)) = 3.83 + 0.92(ln(size_t))$ |
| | Recruit size | $c_o(z') = N(\mu_r, \sigma_r)$; $\mu_r = 0.19$; $\sigma_r = 0.80$ |
| F | Survival | $logit(s(z)) = 0.54 + 0.24(ln(size_t))$ |
| | Flowering | $logit(Pb(z)) = -27.49 + 20.83(ln(size_t)) - 4.04(ln(size_t)^2)$ |
| | Growth | $G(z', z) = N(\mu_s, \sigma_s)$; $\mu_s = 1.62 + 0.16(ln(size_t))$; $\sigma_s = 0.44$ |
| | Seed prod. | $exp(b(z)) = 1.72 + 1.51(ln(size_t))$ |
| | Recruit size | $c_o(z') = N(\mu_r, \sigma_r)$; $\mu_r = 0.15$; $\sigma_r = 0.80$ |
| G | Survival | $logit(s(z)) = -0.19 + 0.30(ln(size_t))$ |
| | Flowering | $logit(Pb(z)) = -32.72 + 23.50(ln(size_t)) - 4.23(ln(size_t)^2)$ |
| | Growth | $G(z', z) = N(\mu_s, \sigma_s)$; $\mu_s = 1.42 + 0.21(ln(size_t))$; $\sigma_s = 0.42$ |
| | Seed prod. | $exp(b(z)) = 0.83 + 2.06(ln(size_t))$ |
| | Recruit size | $c_o(z') = N(\mu_r, \sigma_r)$; $\mu_r = 0.21$; $\sigma_r = 0.77$ |
| H | Survival | $logit(s(z)) = -0.32 + 0.62(ln(size_t))$ |
| | Flowering | $logit(Pb(z)) = -31.04 + 23.96(ln(size_t)) - 4.71(ln(size_t)^2)$ |
| | Growth | $G(z', z) = N(\mu_s, \sigma_s)$; $\mu_s = 1.50 + 0.05(ln(size_t))$; $\sigma_s = 0.43$ |
| | Seed prod. | $exp(b(z)) = 2.53 + 1.59(ln(size_t))$ |
| | Recruit size | $c_o(z') = N(\mu_r, \sigma_r)$; $\mu_r = 0.23$; $\sigma_r = 0.76$ |
| I | Survival | $logit(s(z)) = -0.33 + 0.46(ln(size_t)) - 0.0003(N_t)$ |

| | | |
|---|---|---|
| | Flowering | $logit(Pb(z)) = -41.36 + 33.73(ln(size_t)) - 7.21(ln(size_t)^2) + 0.0008(N_t)$ |
| | Growth | $G(z', z) = N(\mu_s, \sigma_s)$; $\mu_s = 2.19 + 0.14(ln(size_t)) - 0.0005(N_t)$; $\sigma_s = 0.42$ |
| | Seed prod. | $exp(b(z)) = 3.73 + 1.01(ln(size_t))$ |
| | Recruit size | $c_o(z') = N(\mu_r, \sigma_r)$; $\mu_r = 0.19$; $\sigma_r = 0.76$ |
| J | Survival | $logit(s(z)) = 17.59 + 0.05(ln(size_t)) - 0.03(N_t)$ |
| | Flowering | $logit(Pb(z)) = -46.27 + 37.38(ln(size_t)) - 7.83(ln(size_t)^2) + 0.002(N_t)$ |
| | Growth | $G(z', z) = N(\mu_s, \sigma_s)$; $\mu_s = 3.70 + 0.28(ln(size_t)) - 0.004(N_t)$; $\sigma_s = 0.51$ |
| | Seed prod. | $exp(b(z)) = 3.80 + 1.11(ln(size_t))$ |
| | Recruit size | $c_o(z') = N(\mu_r, \sigma_r)$; $\mu_r = 0.22$; $\sigma_r = 0.78$ |
| K | Survival | $logit(s(z)) = -13.85 + 0.20(ln(size_t)) + 0.04(N_t)$ |
| | Flowering | $logit(Pb(z)) = -21.21 + 15.50(ln(size_t)) - 3.55(ln(size_t)^2) + 0.009(N_t)$ |
| | Growth | $G(z', z) = N(\mu_s, \sigma_s)$; $\mu_s = -0.49 + 0.24(ln(size_t)) + 0.006(N_t)$; $\sigma_s = 0.48$ |
| | Seed prod. | $exp(b(z)) = 3.83 + 0.92(ln(size_t))$ |
| | Recruit size | $c_o(z') = N(\mu_r, \sigma_r)$; $\mu_r = 0.19$; $\sigma_r = 0.80$ |
| L | Survival | $logit(s(z)) = 33.06 + 0.24(ln(size_t)) - 0.44(N_t)$ |
| | Flowering | $logit(Pb(z)) = -26.23 + 21.0(ln(size_t)) - 4.01(ln(size_t)^2) - 0.02(N_t)$ |
| | Growth | $G(z', z) = N(\mu_s, \sigma_s)$; $\mu_s = 15.89 + 0.17(ln(size_t)) - 0.19(N_t)$; $\sigma_s = 0.43$ |
| | Seed prod. | $exp(b(z)) = 1.72 + 1.51(ln(size_t))$ |
| | Recruit size | $c_o(z') = N(\mu_r, \sigma_r)$; $\mu_r = 0.15$; $\sigma_r = 0.80$ |
| M | Survival | $logit(s(z)) = 5.39 + 0.37(ln(size_t)) - 0.02(N_t)$ |
| | Flowering | $logit(Pb(z)) = -32.79 + 24.02(ln(size_t)) - 4.34(ln(size_t)^2) - 0.002(N_t)$ |
| | Growth | $G(z', z) = N(\mu_s, \sigma_s)$; $\mu_s = 2.26 + 0.23(ln(size_t)) - 0.003(N_t)$; $\sigma_s = 0.39$ |
| | Seed prod. | $exp(b(z)) = 0.83 + 2.06(ln(size_t))$ |
| | Recruit size | $c_o(z') = N(\mu_r, \sigma_r)$; $\mu_r = 0.21$; $\sigma_r = 0.77$ |
| N | Survival | $logit(s(z)) = 4.30 + 0.87(ln(size_t)) - 0.004(N_t)$ |

| | | |
|---|---|---|
| | Flowering | $logit(Pb(z)) = -28.50 + 25.52(ln(size_t)) - 4.93(ln(size_t)^2) - 0.004(N_t)$ |
| | Growth | $G(z', z) = N(\mu_s, \sigma_s)$; $\mu_s = 2.76 + 0.16(ln(size_t)) - 0.001(N_t)$; $\sigma_s = 0.36$ |
| | Seed prod. | $exp(b(z)) = 2.53 + 1.59(ln(size_t))$ |
| | Recruit size | $c_o(z') = N(\mu_r, \sigma_r)$; $\mu_r = 0.23$; $\sigma_r = 0.77$ |
| S | Survival | $logit(s(z)) = 1.11 + 0.49(ln(size_t)) + 0.005(N_t) - 0.28(T_G)$ |
| | Flowering | $logit(Pb(z)) = -47.12 + 33.70(ln(size_t)) - 7.21(ln(size_t)^2) + 0.0004(N_t) + 0.42(T_G) - 0.23(T_W)$ |
| | Growth | $G(z', z) = N(\mu_s, \sigma_s)$; $\mu_s = 4.12 + 0.14(ln(size_t)) + 0.0009(N_t) - 0.20(T_G)$; $\sigma_s = 0.41$ |
| | Seed prod. | $exp(b(z)) = 3.52+0.90(ln(size_t))+0.0008(N_t)+0.01(T_G)+0.10(T_W)$ |
| | Recruit size | $c_o(z') = N(\mu_r, \sigma_r)$; $\mu_r = -0.38+0.00005(N_t)+0.04(T_G)+0.03(T_W)$; $\sigma_r = 0.76$ |
| T | Survival | $logit(s(z)) = -27.59 + 0.21(ln(size_t)) - 0.0006(N_t) + 1.91(T_G)$ |
| | Flowering | $logit(Pb(z)) = -70.28 + 35.37(ln(size_t)) - 7.45(ln(size_t)^2) - 0.003(N_t) + 1.83(T_G) - 0.95(T_W)$ |
| | Growth | $G(z', z) = N(\mu_s, \sigma_s)$; $\mu_s = -2.88 + 0.27(ln(size_t)) + 0.0002(N_t) + 0.32(T_G)$; $\sigma_s = 0.49$ |
| | Seed prod. | $exp(b(z)) = -12.0 + 0.80(ln(size_t)) - 0.002(N_t) + 1.07(T_G) - 0.94(T_W)$ |
| | Recruit size | $c_o(z') = N(\mu_r, \sigma_r)$; $\mu_r = -0.65 - 0.0007(N_t) + 0.06(T_G) - 0.16(T_W)$; $\sigma_r = 0.77$ |
| U | Survival | $logit(s(z)) = -14.02 + 0.39(ln(size_t)) + 0.006(N_t) + 0.87(T_G)$ |
| | Flowering | $logit(Pb(z)) = -22.83 + 16.0(ln(size_t)) - 3.66(ln(size_t)^2) + 0.002(N_t) + 0.19(T_G) - 1.01(T_W)$ |
| | Growth | $G(z', z) = N(\mu_s, \sigma_s)$; $\mu_s = 0.40 + 0.25(ln(size_t)) - 0.001(N_t) + 0.10(T_G)$; $\sigma_s = 0.46$ |
| | Seed prod. | $exp(b(z)) = -14.33 + 0.59(ln(size_t)) + 0.001(N_t) + 1.18(T_G) - 1.17(T_W)$ |
| | Recruit size | $c_o(z') = N(\mu_r, \sigma_r)$; $\mu_r = -0.56 - 0.00002(N_t)+0.05(T_G)+0.09(T_W)$; $\sigma_r = 0.81$ |
| V | Survival | $logit(s(z)) = -6.78 + 0.24(ln(size_t)) - 0.08(N_t) + 0.69(T_G)$ |

| | | |
|---|---|---|
| | Flowering | $logit(Pb(z)) = -36.50 + 18.80(ln(size_t)) - 3.57(ln(size_t)^2) - 0.02(N_t) + 0.79(T_G) + 0.09(T_W)$ |
| | Growth | $G(z', z) = N(\mu_s, \sigma_s)$; $\mu_s = -1.83 + 0.17(ln(size_t)) + 0.0003(N_t) + 0.23(T_G)$; $\sigma_s = 0.42$ |
| | Seed prod. | $exp(b(z)) = 8.44 + 1.32(ln(size_t)) + 0.003(N_t) - 0.43(T_G) - 0.18(T_W)$ |
| | Recruit size | $c_o(z') = N(\mu_r, \sigma_r)$; $\mu_r = -2.01 + 0.006(N_t) + 0.13(T_G) - 0.01(T_W)$; $\sigma_r = 0.80$ |
| W | Survival | $logit(s(z)) = -32.30 + 0.38(ln(size_t)) + 0.0008(N_t) + 2.19(T_G)$ |
| | Flowering | $logit(Pb(z)) = -311.80 + 18.72(ln(size_t)) - 3.56(ln(size_t)^2) - 0.002(N_t) + 17.93(T_G) - 20.46(T_W)$ |
| | Growth | $G(z', z) = N(\mu_s, \sigma_s)$; $\mu_s = -2.73 + 0.22(ln(size_t)) - 0.0006(N_t) + 0.29(T_G)$; $\sigma_s = 0.39$ |
| | Seed prod. | $exp(b(z)) = -7.07 + 1.47(ln(size_t)) + 0.003(N_t) + 0.63(T_G)$ |
| | Recruit size | $c_o(z') = N(\mu_r, \sigma_r)$; $\mu_r = 1.75 + 0.006(N_t) - 0.10(T_G) + 0.17(T_W)$; $\sigma_r = 0.77$ |
| X | Survival | $logit(s(z)) = -34.59 + 0.88(ln(size_t)) + 0.001(N_t) + 2.26(T_G)$ |
| | Flowering | $logit(Pb(z)) = -43.64 + 28.21(ln(size_t)) - 5.35(ln(size_t)^2) - 0.001(N_t) + 0.57(T_G) + 1.33(T_W)$ |
| | Growth | $G(z', z) = N(\mu_s, \sigma_s)$; $\mu_s = -7.65 + 0.16(ln(size_t)) - 0.0000004(N_t) + 0.61(T_G)$; $\sigma_s = 0.36$ |
| | Seed prod. | $exp(b(z)) = -28.65 + 0.62(ln(size_t)) + 0.0005(N_t) + 2.15(T_G) - 1.64(T_W)$ |
| | Recruit size | $c_o(z') = N(\mu_r, \sigma_r)$; $\mu_r = 0.48 - 0.00004(N_t) - 0.01(T_G) + 0.005(T_W)$; $\sigma_r = 0.77$ |
| AA | Survival | $logit(s(z)) = -0.26 + 0.54(ln(size_t))$ |
| | Flowering | $logit(Pb(z)) = -28.96 + 21.71(ln(size_t)) - 4.14(ln(size_t)^2)$ |
| | Growth | $G(z', z) = N(\mu_s, \sigma_s)$; $\mu_s = 1.48 + 0.10(ln(size_t))$; $\sigma_s = 0.44$ |
| | Seed prod. | $exp(b(z)) = 3.12 + 1.24(ln(size_t))$ |
| | Recruit size | $c_o(z') = N(\mu_r, \sigma_r)$; $\mu_r = 0.23$; $\sigma_r = 0.77$ |
| BB | Survival | $logit(s(z)) = -0.18 + 0.29(ln(size_t))$ |
| | Flowering | $logit(Pb(z)) = -32.70 + 26.93(ln(size_t)) - 5.73(ln(size_t)^2)$ |
| | Growth | $G(z', z) = N(\mu_s, \sigma_s)$; $\mu_s = 1.72 + 0.18(ln(size_t))$; $\sigma_s = 0.51$ |
| | Seed prod. | $exp(b(z)) = 3.45 + 1.17(ln(size_t))$ |
| | Recruit size | $c_o(z') = N(\mu_r, \sigma_r)$; $\mu_r = 0.19$; $\sigma_r = 0.77$ |

| | | |
|---|---|---|
| CC | Survival | $logit(s(z)) = -1.23 + 0.76(ln(size_t))$ |
| | Flowering | $logit(Pb(z)) = -45.81 + 38.00(ln(size_t)) - 8.03(ln(size_t)^2)$ |
| | Growth | $G(z', z) = N(\mu_s, \sigma_s)$; $\mu_s = 1.57 + 0.12(ln(size_t))$; $\sigma_s = 0.44$ |
| | Seed prod. | $exp(b(z)) = 3.21 + 1.22(ln(size_t))$ |
| | Recruit size | $c_o(z') = N(\mu_r, \sigma_r)$; $\mu_r = 0.13$; $\sigma_r = 0.77$ |
| DD | Survival | $logit(s(z)) = 2.43 - 0.23(ln(size_t))$ |
| | Flowering | $logit(Pb(z)) = -86.83 + 71.52(ln(size_t)) - 14.65(ln(size_t)^2)$ |
| | Growth | $G(z', z) = N(\mu_s, \sigma_s)$; $\mu_s = 1.93 + 0.25(ln(size_t))$; $\sigma_s = 0.56$ |
| | Seed prod. | $exp(b(z)) = 6.01 + 0.34(ln(size_t))$ |
| | Recruit size | $c_o(z') = N(\mu_r, \sigma_r)$; $\mu_r = 0.30$; $\sigma_r = 0.69$ |
| EE | Survival | $logit(s(z)) = 0.65 + 0.15(ln(size_t))$ |
| | Flowering | $logit(Pb(z)) = -24.35 + 17.91(ln(size_t)) - 3.49(ln(size_t)^2)$ |
| | Growth | $G(z', z) = N(\mu_s, \sigma_s)$; $\mu_s = 1.85 + 0.20(ln(size_t))$; $\sigma_s = 0.52$ |
| | Seed prod. | $exp(b(z)) = 2.51 + 1.54(ln(size_t))$ |
| | Recruit size | $c_o(z') = N(\mu_r, \sigma_r)$; $\mu_r = 0.09$; $\sigma_r = 0.86$ |
| FF | Survival | $logit(s(z)) = 1.50 - 0.32(ln(size_t))$ |
| | Flowering | $logit(Pb(z)) = -70.62 + 54.30(ln(size_t)) - 10.46(ln(size_t)^2)$ |
| | Growth | $G(z', z) = N(\mu_s, \sigma_s)$; $\mu_s = 1.65 + 0.21(ln(size_t))$; $\sigma_s = 0.46$ |
| | Seed prod. | $exp(b(z)) = 1.91 + 1.39(ln(size_t))$ |
| | Recruit size | $c_o(z') = N(\mu_r, \sigma_r)$; $\mu_r = 0.10$; $\sigma_r = 0.92$ |
| GG | Survival | $logit(s(z)) = 0.99 + 0.12(ln(size_t))$ |
| | Flowering | $logit(Pb(z)) = -50.32 + 36.50(ln(size_t)) - 6.54(ln(size_t)^2)$ |
| | Growth | $G(z', z) = N(\mu_s, \sigma_s)$; $\mu_s = 1.48 + 0.28(ln(size_t))$; $\sigma_s = 0.38$ |
| | Seed prod. | $exp(b(z)) = 3.06 + 1.22(ln(size_t))$ |
| | Recruit size | $c_o(z') = N(\mu_r, \sigma_r)$; $\mu_r = 0.18$; $\sigma_r = 0.81$ |
| HH | Survival | $logit(s(z)) = 0.74 + 0.33(ln(size_t))$ |
| | Flowering | $logit(Pb(z)) = -50.91 + 40.17(ln(size_t)) - 7.81(ln(size_t)^2)$ |
| | Growth | $G(z', z) = N(\mu_s, \sigma_s)$; $\mu_s = 1.62 + 0.18(ln(size_t))$; $\sigma_s = 0.39$ |
| | Seed prod. | $exp(b(z)) = 4.20 + 0.97(ln(size_t))$ |
| | Recruit size | $c_o(z') = N(\mu_r, \sigma_r)$; $\mu_r = 0.24$; $\sigma_r = 0.76$ |
| II | Survival | $logit(s(z)) = -0.26 + 0.06(ln(size_t))$ |
| | Flowering | $logit(Pb(z)) = -41.11 + 36.347(ln(size_t)) - 8.31(ln(size_t)^2)$ |
| | Growth | $G(z', z) = N(\mu_s, \sigma_s)$; $\mu_s = 1.67 + 0.16(ln(size_t))$; $\sigma_s = 0.38$ |
| | Seed prod. | $exp(b(z)) = 4.92 + 0.41(ln(size_t))$ |

| | | |
|---|---|---|
| | Recruit size | $c_o(z') = N(\mu_r, \sigma_r)$; $\mu_r = 0.22$; $\sigma_r = 0.71$ |
| JJ | Survival | $logit(s(z)) = 0.07 + 0.16(ln(size_t))$ |
| | Flowering | $logit(Pb(z)) = -24.83 + 19.82(ln(size_t)) - 4.20(ln(size_t)^2)$ |
| | Growth | $G(z', z) = N(\mu_s, \sigma_s)$; $\mu_s = 1.63 + 0.32(ln(size_t))$; $\sigma_s = 0.44$ |
| | Seed prod. | $exp(b(z)) = 4.63 + 0.66(ln(size_t))$ |
| | Recruit size | $c_o(z') = N(\mu_r, \sigma_r)$; $\mu_r = 0.09$; $\sigma_r = 0.82$ |
| KK | Survival | $logit(s(z)) = -0.26 + 0.25(ln(size_t))$ |
| | Flowering | $logit(Pb(z)) = -27.90 + 27.66(ln(size_t)) - 7.00(ln(size_t)^2)$ |
| | Growth | $G(z', z) = N(\mu_s, \sigma_s)$; $\mu_s = 1.56 + 0.30(ln(size_t))$; $\sigma_s = 0.41$ |
| | Seed prod. | $exp(b(z)) = 5.72 - 0.09(ln(size_t))$ |
| | Recruit size | $c_o(z') = N(\mu_r, \sigma_r)$; $\mu_r = 0.80$; $\sigma_r = 0.79$ |
| LL | Survival | $logit(s(z)) = -0.22 + 0.77(ln(size_t))$ |
| | Flowering | $logit(Pb(z)) = -147.87 + 121.01(ln(size_t)) - 24.82(ln(size_t)^2)$ |
| | Growth | $G(z', z) = N(\mu_s, \sigma_s)$; $\mu_s = 1.61 + 0.10(ln(size_t))$; $\sigma_s = 0.39$ |
| | Seed prod. | $exp(b(z)) = 1.98 + 1.57(ln(size_t))$ |
| | Recruit size | $c_o(z') = N(\mu_r, \sigma_r)$; $\mu_r = 0.16$; $\sigma_r = 0.68$ |
| MM | Survival | $logit(s(z)) = -1.13 + 0.60(ln(size_t))$ |
| | Flowering | $logit(Pb(z)) = -45.56 + 39.31(ln(size_t)) - 8.64(ln(size_t)^2)$ |
| | Growth | $G(z', z) = N(\mu_s, \sigma_s)$; $\mu_s = 1.34 + 0.14(ln(size_t))$; $\sigma_s = 0.40$ |
| | Seed prod. | $exp(b(z)) = 0.71 + 2.00(ln(size_t))$ |
| | Recruit size | $c_o(z') = N(\mu_r, \sigma_r)$; $\mu_r = 0.32$; $\sigma_r = 0.64$ |
| NN | Survival | $logit(s(z)) = -1.74 + 1.39(ln(size_t))$ |
| | Flowering | $logit(Pb(z)) = -27.89 + 20.50(ln(size_t)) - 3.99(ln(size_t)^2)$ |
| | Growth | $G(z', z) = N(\mu_s, \sigma_s)$; $\mu_s = 1.17 + 0.13(ln(size_t))$; $\sigma_s = 0.33$ |
| | Seed prod. | $exp(b(z)) = 5.49 + 0.14(ln(size_t))$ |
| | Recruit size | $c_o(z') = N(\mu_r, \sigma_r)$; $\mu_r = 0.24$; $\sigma_r = 0.78$ |

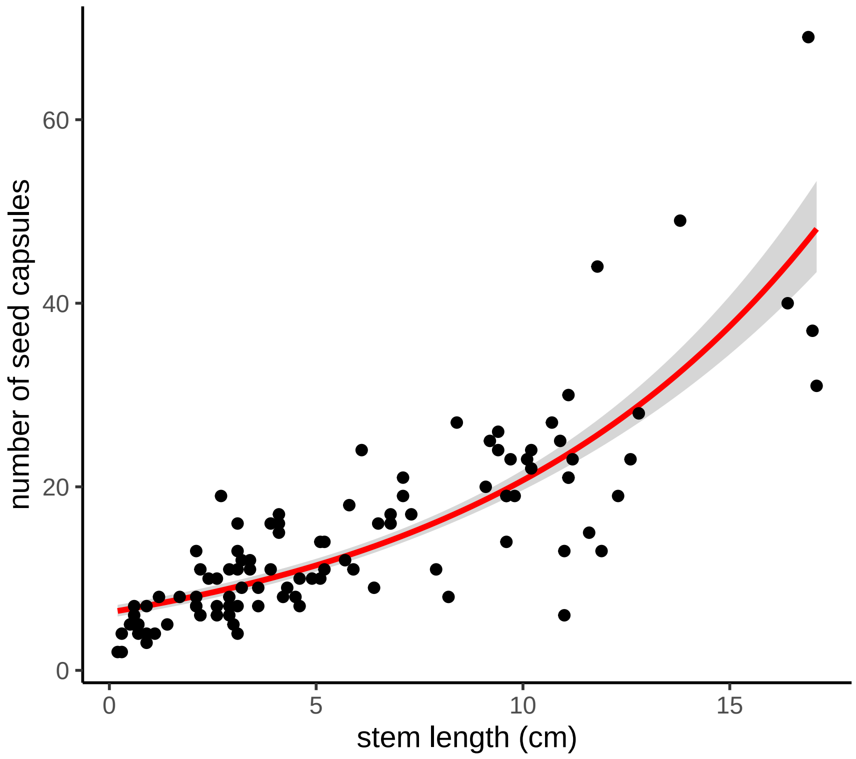


Figure S2: As the stem length of an *Oenothera coloradensis* flowering individual increases, the number of capsules it produces increases as well. The red line shows the fit from a Poisson generalized linear model, and the grey ribbon shows the 95% confidence interval around the fitted relationship. Model equation: Number of capsules = $e^{(1.843+0.119\text{x}S))}$, where $S$ is stem length in cm (pseudo R-squared = 0.42, $P = < 0.01$, Residual deviance = 186.98, df = 104).

Table S3: Comparison of vital rate models with and without density dependence. The "DI" and "DD" rows contain AIC values for each vital rate model in each subpopulation for models that are density-independent (DI) and density-dependent (DD). The difference between the AIC of DI and DD models is shown in the ΔAIC column. Bold text indicates that the |Δ|AIC value is > 3, which means that including a term for density dependence substantially changed that vital rate model. A positive |Δ|AIC indicates that including density dependence improved the model, while a negative value indicates that including density dependence made model fit worse. AIC values for DI and DD values can be found in Table S3.

| Vital Rate Model | | Subpopulation | | | | | |
|---|---|---|---|---|---|---|---|
| | | Crow Creek | Diamond Creek | Unnamed Creek | HQ5 | HQ3 | Meadow |
| Survival | DI | 776.58 | 1012.68 | 2684.34 | 3242.63 | 716.66 | 166.13 |
| | DD | 757.84 | 905.39 | 26848.74 | 2922.91 | 637.84 | 166.83 |
| | ΔAIC | **18.74** | **107.28** | -0.41 | **320.33** | **78.82** | -0.70 |
| Growth | DI | 510.34 | 953.29 | 1098.95 | 1570.93 | 300.18 | 116.54 |
| | DD | 506.61 | 931.15 | 1068.14 | 1112.78 | 269.73 | 113.88 |
| | ΔAIC | **3.73** | **22.15** | **30.811** | **458.15** | **30.45** | 2.66 |
| Flowering | DI | 371.68 | 523.30 | 1087.93 | 538.52 | 191.46 | 104.24 |
| | DD | 373.31 | 523.74 | 1087.48 | 483.99 | 193.22 | 106.96 |
| | ΔAIC | -1.63 | -0.44 | 0.45 | **54.52** | -1.76 | -1.72 |
| Seed production | DI | 842.00 | 1580.85 | 2815.89 | 1423.02 | 598.75 | 280.09 |
| | DD | 835.59 | 1566.83 | 2817.19 | 1419.32 | 594.63 | 281.45 |
| | ΔAIC | **6.41** | **14.02** | -1.29 | **3.71** | **4.12** | -1.35 |
| Recruit size | DI | 921.31 | 1028.23 | 3378.43 | 4629.87 | 967.83 | 173.03 |
| | DD | 923.24 | 1026.63 | 3380.53 | 4631.84 | 969.06 | 175.02 |
| | ΔAIC | -1.93 | 1.61 | -1.93 | -1.97 | -1.23 | -1.99 |
